# Emergence of trait variability through the lens of nitrogen assimilation in *Prochlorococcus*

**DOI:** 10.1101/383927

**Authors:** Paul M. Berube, Anna Rasmussen, Rogier Braakman, Ramunas Stepanauskas, Sallie W. Chisholm

## Abstract

Intraspecific trait variability has important consequences for the function and stability of marine ecosystems. The marine cyanobacterium *Prochlorococcus* is a useful model system for understanding how trait variability emerges within microbial species: Its functional diversity is overlaid on measurable environmental gradients, providing a powerful lens into large-scale evolutionary processes. Here we examine variation in the ability to use nitrate across hundreds of *Prochlorococcus* genomes to better understand the modes of evolution influencing the allocation of ecologically important functions within microbial species. We find that nitrate assimilation genes are absent in basal lineages of *Prochlorococcus* but occur at an intermediate frequency that is randomly distributed within recently emerged clades. The distribution of nitrate assimilation genes within clades appears largely governed by vertical inheritance, stochastic gene loss, and homologous recombination among closely related cells. By mapping this process onto a model of *Prochlorococcus’* macroevolution, we propose that niche-constructing adaptive radiations and subsequent niche partitioning set the stage for loss of nitrate assimilation genes from basal lineages as they specialized to lower light levels. Retention of these genes in recently emerged lineages has likely been facilitated by selection as they sequentially partitioned into niches where nitrate assimilation conferred a fitness benefit.

## INTRODUCTION

*Prochlorococcus* and its closest relative, *Synechococcus*, are non-nitrogen-fixing cyanobacteria whose common ancestor emerged approximately 823–644 Mya (Sánchez-Baracaldo et al. 2014). They are among the most abundant photosynthetic organisms on the planet and are estimated to be jointly responsible for 25% of ocean net primary productivity (Flombaum et al. 2013). While *Synechococcus* has a broader geographic range, *Prochlorococcus* is primarily restricted to the tropical and subtropical open ocean where nutrients are scarce. Genomic signatures of adaptation to these highly oligotrophic environments are found across *Prochlorococcus* genomes (Rocap et al. 2003; Martiny et al. 2006; Kettler et al. 2007) and reveal the selection pressures operating on populations in the wild (Malmstrom et al. 2010; Coleman and Chisholm 2010; Rusch et al. 2010).

Access to nitrogen – often the proximal limiting nutrient for phytoplankton growth across the global ocean (Tyrrell 1999) – is a nearly ubiquitous challenge facing *Prochlorococcus*. All *Prochlorococcus* can assimilate ammonium, but the remainder of nitrogen uptake pathways – for amino acids, urea, cyanate, nitrite, and nitrate – are encoded by flexible genes that are found in some, but not all, *Prochlorococcus*. Nitrate assimilation is of particular interest because nitrate is abundant at the base of the sunlit euphotic zone and can fuel productivity in nitrogen limited surface waters during the vertical advection of this deeper water (Dugdale and Goering 1967; Eppley et al. 1979; Johnson et al. 2010). Nearly all observed marine *Synechococcus* have the genes encoding the transporters, reductases, and molybdopterin cofactor biosynthesis proteins required for nitrate assimilation (Scanlan et al. 2009; Ahlgren and Rocap 2006). Most *Prochlorococcus* lack this genetic repertoire, possibly because nitrate assimilation is a costly pathway; as the most oxidized inorganic nitrogen source in marine systems, the assimilation of nitrate requires 10 electrons compared with 2 electrons for ammonium assimilation (GarcÍa-Fernández et al. 2004). Further, cells must allocate a significant amount of iron to this pathway in the form of ferredoxin as well as multiple iron-containing prosthetic groups that are associated with the reductases (Flores et al. 2005). The additional demand for reducing power, protein biosynthesis, and limiting trace metals likely represents a sizable cost for these oligotrophic cyanobacteria (GarcÍa-Fernández et al. 2004).

*Prochlorococcus* has diverged into multiple clades, many with adaptations that can be mapped onto gradients of key environmental variables (Biller et al. 2015). Among the dozens of cultured *Prochlorococcus* strains sequenced from these clades, genes encoding nitrate assimilation proteins have only been observed in a few strains belonging to the high-light adapted HLII clade and the low-light adapted LLI clade (Berube et al. 2015). In the open ocean, the distribution of *Prochlorococcus* with the potential to assimilate nitrate shows distinct seasonal patterns, reaching a peak in abundance in the summer when the solar energy supply is greater and overall nitrogen concentrations are lower (Berube et al. 2016).

Within the HLII clade – often the most abundant *Prochlorococcus* clade in subtropical gyres by an order of magnitude (Malmstrom et al. 2010) – the frequency of cells that are capable of assimilating nitrate (up to 20-50%) is positively correlated with decreased nitrogen availability in surface waters where they dominate. This has suggested that cells with access to a wide pool of nitrogen sources and a ready supply of energy are likely at a selective advantage when nitrogen is limiting (Berube et al. 2016). For the LLI clade, the trade-offs may be slightly more complex. All previously described *Prochlorococcus* in the LLI clade can assimilate nitrite, but only a fraction can also assimilate the more oxidized nitrate (Berube et al. 2015). These cells are found in the mid-euphotic zone and near the nitracline where inorganic nitrogen concentrations begin to rise with increasing depth. The depth distribution of these low-light adapted cells is associated with a peak in nitrite concentration (Berube et al. 2016). In stratified marine systems, the depth of this nitrite maximum layer is commonly correlated with the nitracline and the depth at which photosynthetically active radiation is attenuated to ~1% of surface irradiance (Lomas and Lipschultz 2006). At this inflection point in the water column, cells have access to inorganic nitrogen sources of varying redox state (NO_3_^-^, NO_2_ ^-^, and NH_4_ ^+^), but must also compete with ammonia and nitrite oxidizing microorganisms for access to reduced and thus more easily assimilated nitrogen sources (Berube et al. 2016). Frequency-dependent selection processes (Cordero and Polz 2014) have a potential role in selecting for the optimal distribution of nitrogen assimilation traits across these low-light adapted *Prochlorococcus* populations.

While we have gained general insights into the tradeoffs governing the overall abundance of *Prochlorococcus* cells capable of nitrate assimilation in the wild, genomic data have thus far painted an incomplete picture of the evolution of this trait at the genomic and sequence levels. Although the gene order and genomic location of the nitrate assimilation gene cluster among HLII clade genomes is highly conserved (Martiny et al. 2009; Berube et al. 2015), there is evidence that one genome acquired duplicate copies of these genes via phage-mediated gene transfer (Berube et al. 2015). Comparative genomics of a small number of cultured strains suggested that *Prochlorococcus* lost the nitrate assimilation genes following their divergence from *Synechococcus* and then regained them early in the emergence of the LLI and HLII clades (Berube et al. 2015). Examination of GC content, gene synteny, and local genomic architecture, however, suggested that nitrate assimilation genes may have descended vertically in *Prochlorococcus* (Berube et al. 2015). Overall, the relative roles of vertical descent, gene duplication, genome streamlining, and horizontal gene transfer in mediating the evolution of these genes is thus ambiguous – especially given the limited number of cultivated representatives among many *Prochlorococcus* clades.

In this study, we examined a genomics data set consisting of 486 *Prochlorococcus* and 59*Synechococcus* genomes from both cultivated isolates and wild single cells to better constrain the relative impacts of different evolutionary forces on the diversification of nitrate assimilation genes in *Prochlorococcus*. Drawing on our understanding of the physiology and ecology of major phylogenetic groups of *Prochlorococcus*, we also explore how evolution gives rise to a heterogeneous distribution of the nitrate assimilation trait across the *Prochlorococcus* genus.

## RESULTS AND DISCUSSION

### Nitrate assimilation genes are found within a distinct subset of *Prochlorococcus* clades

To constrain the higher order relationship between nitrate assimilation and major clades of *Prochlorococcus*, we first assessed the distribution of genes involved in nitrate and nitrite assimilation across a core marker gene phylogeny for *Prochlorococcus* and *Synechococcus* that includes both isolate and single cell genomes (Fig. 1). Given the inherent incompleteness of single cell genome assemblies, we also developed a PCR assay (Supplementary Methods) to screen amplified single cell DNA for the presence of *narB* (nitrate reductase), a key gene distinguishing the complete pathway for nitrate assimilation from the downstream half for nitrite assimilation alone. We found that while genes for nitrite assimilation are distributed across the tree of *Prochlorococcus*, thus enabling the assimilation of the more reduced nitrite, genes for the full pathway of nitrate assimilation are restricted to particular clades (Fig. 1). Among the high-light adapted *Prochlorococcus*, *narB* was found in the HLI, HLII, and HLVI clades (Fig. 1). It also appears rare for high-light adapted *Prochlorococcus* genomes to encode only the downstream half of the nitrate assimilation pathway – the reduction of nitrite to ammonium – instead of the complete pathway. Among the polyphyletic group of low-light adapted *Prochlorococcus*, *narB* was exclusively found in the LLI clade (Fig. 1). We therefore conclude that the complete pathway for nitrate assimilation is essentially restricted to clades that emerged more recently in the evolution of *Prochlorococcus.*

**Fig. 1.**
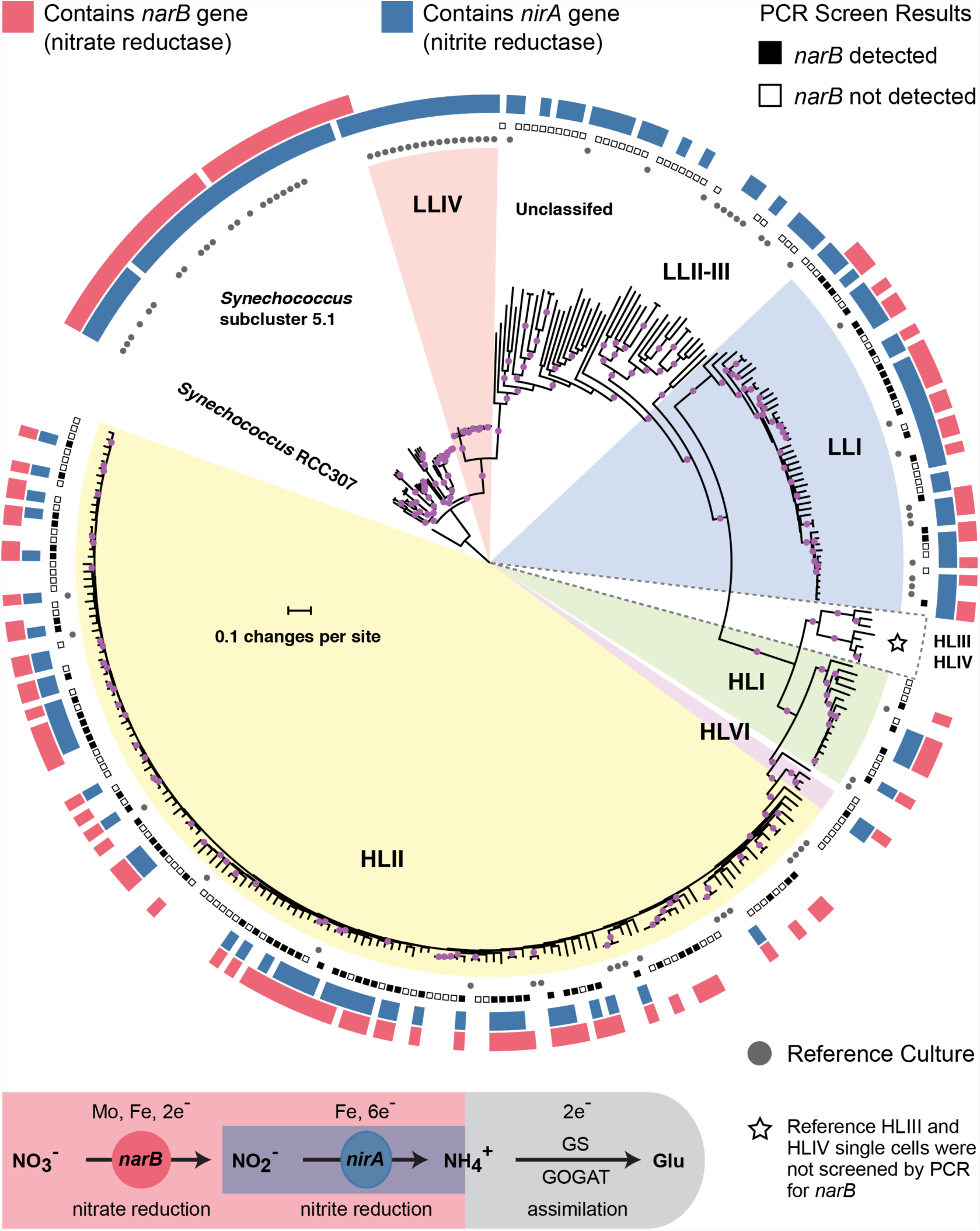
Distributions of the nitrate reductase (*narB*) and nitrite reductase (*nirA*) genes across a core marker gene phylogeny of 329 *Prochlorococcus* and *Synechococcus* genomes. Selected *Prochlorococcus* clades are highlighted as pie slices. Red bars indicate genomes with the potential to use nitrate based on the presence/absence of a *narB* gene. Blue bars indicate genomes with an annotated *nirA* gene in the genome assembly. Reference culture genomes are indicated by gray circles and the results of the PCR screen for *narB* are indicated by filled (present) and open (absent) squares. Single cells belonging to the HLIII and HLIV clades of *Prochlorococcus* were not screened by PCR and are only included as additional reference genomes. The nucleotide phylogeny is based on a concatenated alignment of 37 marker genes in the PhyloSift software package and inferred using maximum likelihood in RAxML (GTRCAT model) with automatic bootstopping criteria (250 replicate trees). Filled purple circles on branches indicate that the associated genomes clustered together in at least 75% of trees.

The apparent absence of *narB* in some *Prochlorococcus* clades is consistent with expectations based on tradeoffs (e.g. energetic and trace metal requirements) that are inferred to govern nitrate assimilation. The HLIII and HLIV clades, which lack this trait in available single cell genome assemblies (Malmstrom et al. 2013), are generally characterized by adaptations to iron limitation (Rusch et al. 2010; Malmstrom et al. 2013) and may thus be under selective pressure to dispense with the nitrate assimilation pathway due to its high iron requirement (Malmstrom et al. 2013). While genes for the downstream half of the nitrate assimilation pathway – encoding the NirA, FocA, and NirX proteins for the transport and reduction of nitrite to ammonium – are present in all low-light adapted clades of *Prochlorococcus* in our data set, *narB* is exclusively associated with the LLI clade (Fig. 1). Cells belonging to the LLI clade dominate at shallower depths than other low-light adapted cells and moreover expand into the surface mixed layer during winter months (Zinser et al. 2007; Malmstrom et al. 2010). Thus, among low-light adapted *Prochlorococcus*, the LLI clade experiences higher irradiance levels which would facilitate the generation of required reducing power equivalents to support the reduction of nitrate. Further, the average genome recovery of single cell genome assemblies belonging to the LLI clade matched the proportion of these assemblies with an annotated *nirA* gene (76% average genome recovery; 76% containing *nirA*), indicating that the ability to assimilate the more reduced nitrite is a core trait for cells belonging to the LLI clade. We suspect that this may be a selective advantage for *Prochlorococcus* that live in close proximity to elevated concentrations of nitrite (Berube et al. 2016) that occur in the ubiquitous primary nitrite maximum (Lomas and Lipschultz 2006).

### Nitrate assimilation genes do not co-vary with other flexible genes

Identifying traits that are under- or over-represented in *Prochlorococcus* genomes that harbor nitrate assimilation genes could shed additional light on other selection pressures similarly operating on cells with this trait and how cells in these clades balance the tradeoffs of the nitrate assimilation pathway. To explore this, we compared the flexible gene content of single cell *Prochlorococcus* genomes that possess *narB* with those that lack *narB* using gene enrichment analysis. No flexible genes other than those involved in nitrate assimilation were found to be over- or under-represented in genomes containing *narB* (hypergeometric test with Benjamini and Hochberg correction; p < 0.05) (Fig. 2 and Supplementary Fig. 1). Although there appears to be a selective advantage to carrying nitrate assimilation genes under nitrogen limiting conditions (Berube et al. 2016), cells capable of using nitrate are no more likely to carry additional accessory nitrogen assimilation pathways such as those for cyanate, urea, or amino acid assimilation. Overall, it appears that the nitrate assimilation genes have evolved independently of other flexible traits in *Prochlorococcus*.

**Fig. 2.**
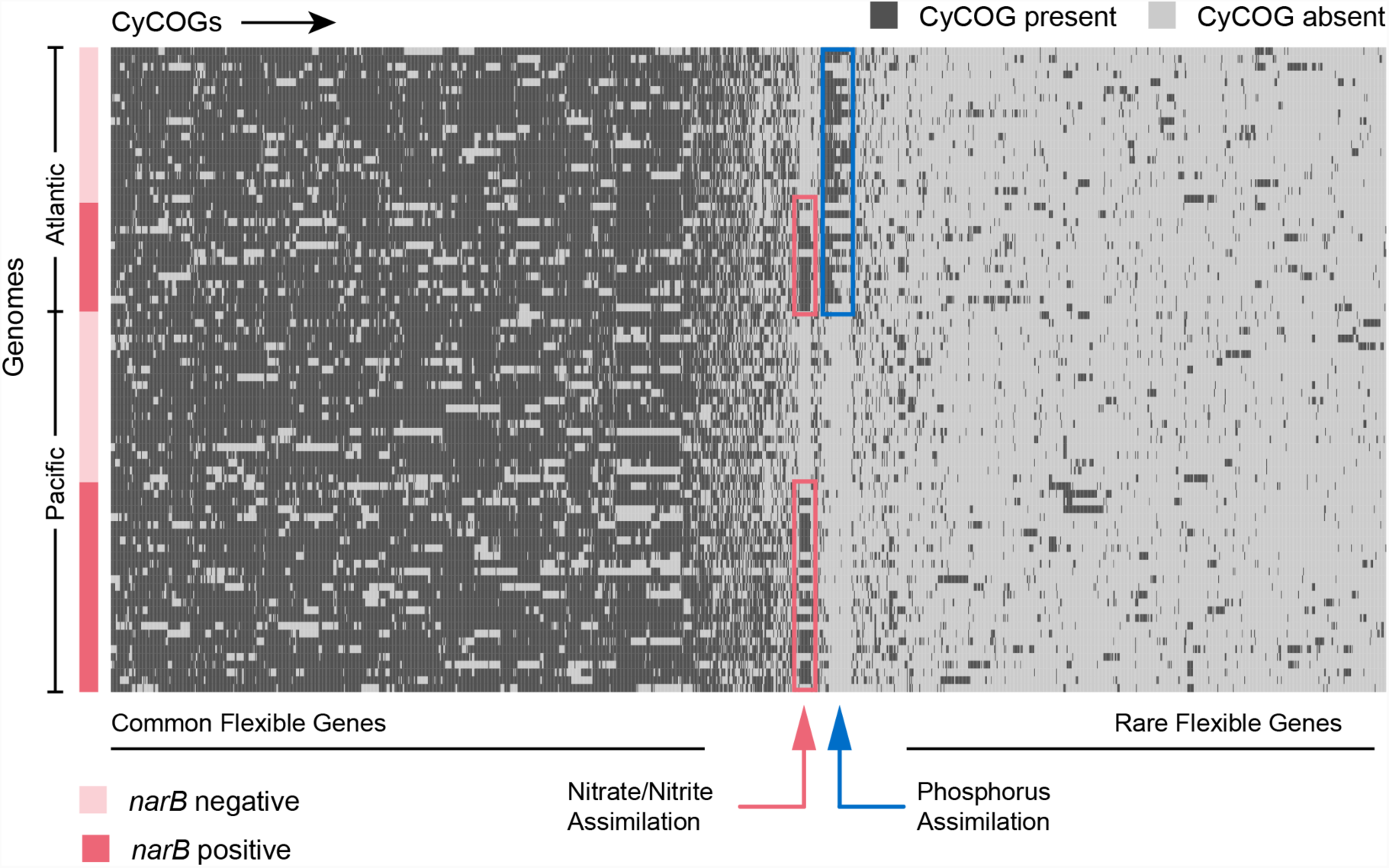
Hierarchical clustering of presence and absence distributions for flexible CyCOGs found in 83 *Prochlorococcus* HLII single cell genomes with genome recoveries of at least 75% (median 90%). Genomes are sorted by Atlantic and Pacific Oceans and by the presence/absence of the *narB* gene – a marker for the capacity to assimilate nitrate. Other than genes in the nitrate assimilation gene cluster (green box), no CyCOGs were over- or under-represented among the flexible genes in genomes containing *narB*. Genomes from the Atlantic, regardless of whether or not they contained the *narB* marker gene, were enriched in phosphorus assimilation genes (blue box).

### Nitrate assimilation genes are rarely acquired by *Prochlorococcus* through non-homologous recombination mechanisms

Trait absence in basal lineages but presence in recently emerged lineages suggests a possible role for horizontal gene transfer: *Prochlorococcus* may have lost the nitrate assimilation genes early after its divergence from *Synechococcus* and reacquired them later through horizontal gene transfer mechanisms (Berube et al. 2015). Other evidence, however, namely the close relationship between gene and whole genome GC contents as well as conservation in the location and gene order of the nitrate assimilation gene cluster, suggests that vertical descent has been an important factor in shaping these genome features (Martiny et al. 2009; Berube et al. 2015). To further assess the role of horizontal acquisition in the evolution of the nitrate assimilation trait in *Prochlorococcus*, we first examined in greater depth the patterns of diversity of this trait within clades.

We compared the phylogenies of individual proteins in both the upstream and downstream halves of the nitrate assimilation pathway with those of core marker proteins (primarily ribosomal proteins) that we assume are vertically inherited. Phylogenies for the nitrate reductase (NarB) and a molybdopterin biosynthesis protein (MoaA) were generally congruent with the core protein phylogeny at the clade level, arguing against frequent horizontal gene transfer between clades (Fig. 3a,c-d). The nitrate transporter (NapA) protein phylogeny (Fig. 3b) exhibited a branching pattern that grouped LLI *Prochlorococcus* with *Synechococcus*, but the corresponding gene phylogeny of *napA* was, however, congruent with the core gene phylogeny (Supplementary Fig. 2). Regardless, both HL and LLI *Prochlorococcus* strains possessed proteins in the upstream half of the nitrate assimilation pathway that were monophyletic and we conclude that recent gene transfer between clades has been minimal or absent.

**Fig. 3.**
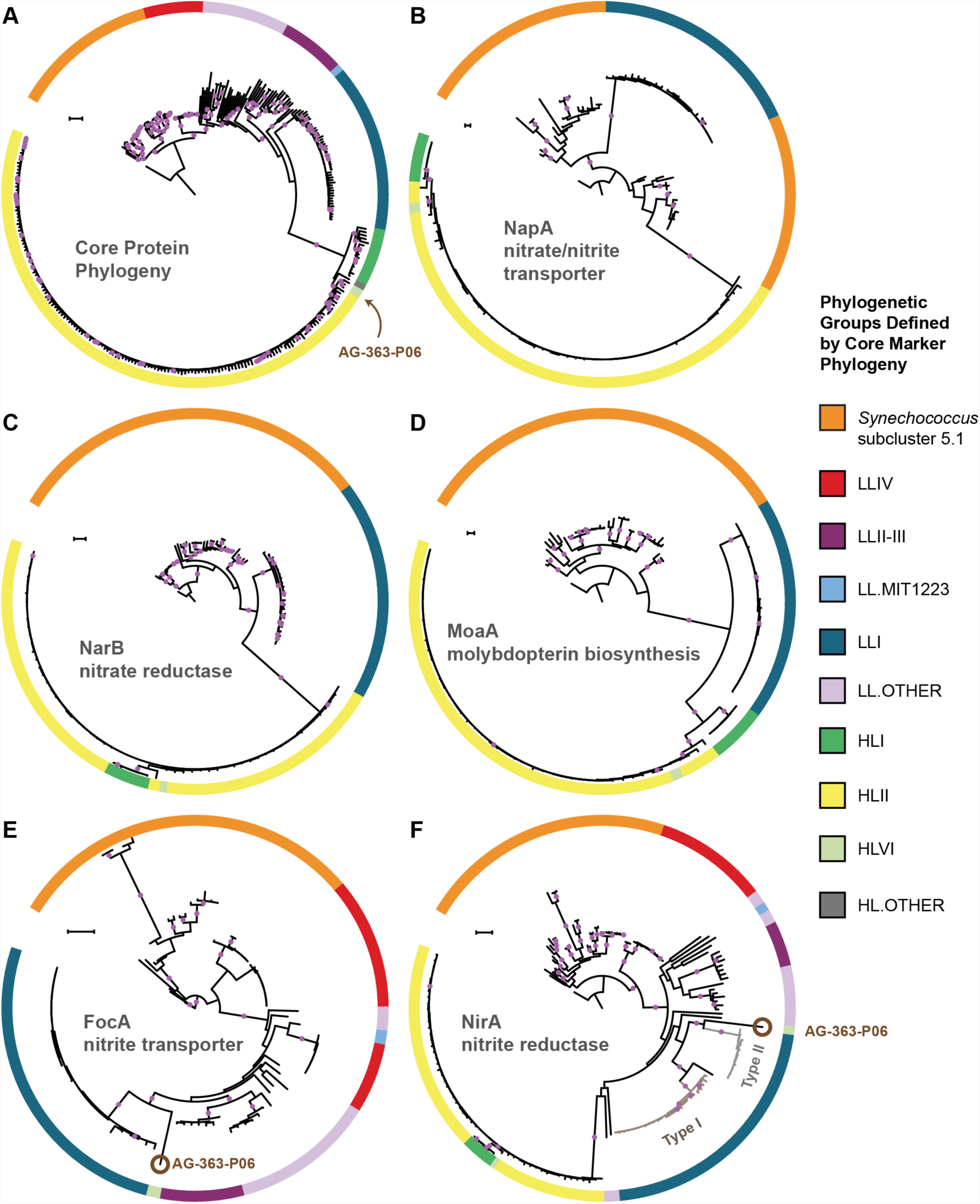
The core marker protein phylogeny of *Prochlorococcus* and *Synechococcus* (A) in comparison to phylogenies for the nitrate/nitrite transporter, NapA (B), the nitrate reductase, NarB (C), the molybdopterin biosynthesis protein, MoaA (D), the nitrite transporter, FocA (E), and the nitrite reductase, NirA (F). Interclade horizontal gene transfer is minimal for genes encoding proteins in the upstream half of the nitrate assimilation pathway (B-D) since clades defined by the core phylogeny (A) remain separate. Horizontal gene transfer is observed in a few instances for genes encoding proteins in the downstream half of the nitrate assimilation pathway (E-F). One single cell (AG-363-P06; brown circle) from the high-light adapted HLVI clade possesses FocA and NirA proteins most similar to those from low-light adapted *Prochlorococcus*. *Prochlorococcus* belonging to the LLI clade possess two types of NirA as indicated by well supported phylogenetic divergence of NirA among LLI cells. Filled purple circles on branches indicate that the associated taxa clustered together in at least 75% of trees. Scale bars are 0.1 changes per site.

Proteins involved in the downstream half of the pathway – responsible for the transport and reduction of nitrite and encoded by a functional cassette with 3 genes (*focA, nirA*, and *nirX*) – had phylogenies that were also largely consistent with the core protein phylogeny (Figs. 3a and 3e-f). But, we noticed one exception: The high-light adapted AG-363-P06 single cell (HLVI clade) possesses genes encoding a nitrite transporter (FocA) and a nitrite reductase (NirA) that are similar to low-light adapted *Prochlorococcus* (Fig. 3e-f). Among the single cells in our data set, this is the only genome belonging to a high-light adapted clade that contains the *focA* gene. This suggests that this HLVI *Prochlorococcus* strain recently acquired the nitrite assimilation cassette from a low-light adapted *Prochlorococcus*.

We next examined closed genomes and individual contigs from single cells to look for aberrations in the location and gene order of the nitrate assimilation gene cluster that might suggest gene acquisition through non-homologous recombination mechanisms. Within the HLII clade, there were 28 genome assemblies containing the complete nitrate assimilation gene cluster with sufficient sequence data extending into adjacent genomic regions on a single contig. In 27 of them, the nitrate assimilation genes were found in a core syntenic genomic region between a conserved gene encoding a sodium-dependent symporter (CyCOG_60001297) and *polA*, encoding DNA polymerase I (Fig. 4 and Supplementary Fig. 3). The remaining HLII genome, MIT0604, is the only documented member of the HLII clade with duplicate copies of the nitrate assimilation gene cluster located in separate genomic islands (Berube et al. 2015).

**Fig. 4.**
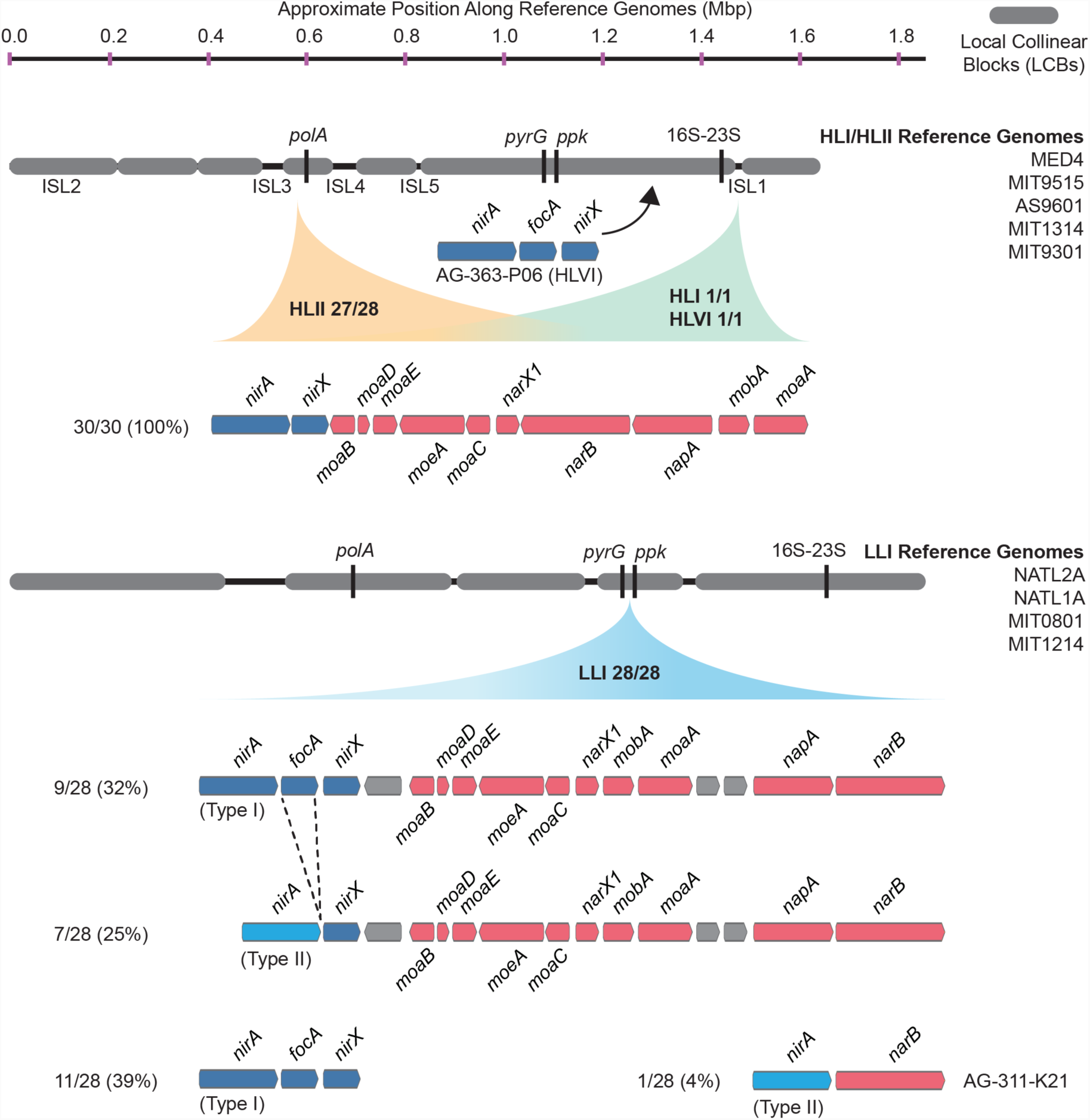
The gene order and genomic location of the nitrate assimilation gene cluster in high-light and low-light adapted *Prochlorococcus* genomes. The location of the nitrate and nitrite assimilation genes is shown relative to conserved core marker genes (black bars). The proportion of genomes in each clade with the indicated location is shown next to the clade names. The percentage of genomes in each group with a specific gene content and order is shown to the left of each gene order plot.

In two single cell genomes belonging to the HLI and HLVI clades, the complete nitrate assimilation gene cluster is found in a different genomic region than in cells belonging to the HLII clade (Fig. 4 and Supplementary Fig. 4). Thus, it appears that the genomic position of these genes is largely conserved within individual high-light adapted clades but can differ between high-light adapted clades. Regardless of their genomic location, gene order was conserved in all genomes belonging to this monophyletic group of high-light adapted clades (Fig. 4). The cell belonging to the HLVI clade (AG-363-P06) that only has the downstream half of the pathway – which we argue above was acquired from low-light adapted *Prochlorococcus* (Fig. 3) – encodes these genes in an entirely different region located upstream of the ribosomal RNA operon (Fig. 4). In this one case, it is likely that gene acquisition was driven by non-homologous recombination mechanisms.

Among 28 LLI clade genomes, we found that the nitrate and nitrite assimilation genes were always located in a core syntenic region between the pyrimidine biosynthesis gene, *pyrG*, and the polyphosphate kinase gene, *ppk* (Fig. 4 and Supplementary Fig. 5), which corresponds with the position of these genes in *Synechococcus* (Rocap et al. 2003; Martiny et al. 2009; Berube et al. 2015), strengthening an argument for vertical inheritance of these genes during the evolution of low-light adapted *Prochlorococcus*. Regardless of whether the LLI clade genomes encode the complete nitrate assimilation pathway or only the downstream half of the pathway (Fig. 4), the order of the genes is conserved (Fig. 4) with one exception: In the AG-311-K21 single cell there was an apparent deletion of the transporters and molybdopterin biosynthesis genes, leaving only the *nirA* and *narB* reductase genes (Fig. 4). Given that both the *pyrG* and *ppk* genes were present in their expected locations in this genome, this is probably a genuine deletion and may reflect initial stages in the loss of the nitrate assimilation pathway in this LLI clade cell. Note, however, that this single cell genome assembly is 46% complete and thus we cannot rule out rearrangement of the missing genes to unassembled regions of the genome. Finally, there is a general pattern in which a small number of LLI genomes lack the nitrite transporter gene, *focA*, and possess a distinct version (Type II) of the nitrite reductase gene, *nirA* (Figs. 3 and 4). These divergent enzymes could be a result of fine-scale niche partitioning, and biochemical studies are warranted to understand their potential functional and ecological significance.

### Homologous recombination shapes the underlying diversity of nitrate assimilation genes in *Prochlorococcus*

Thus far, our analysis has provided little evidence for acquisition of nitrate assimilation genes by *Prochlorococcus* through non-homologous recombination-based mechanisms. Homologous recombination between closely related cells, however, can facilitate the gain and loss of genes through recombination in core genomic regions that flank the genes in question (Apagyi et al. 2018; Oliveira et al. 2017) and can further act to limit the genetic divergence of shared loci (Andam et al. 2010; Andam and Gogarten 2011; Rosen et al. 2015). To explore the potential role of this process, we evaluated the relative influence of recombination and mutation on shaping the diversity of genomic regions containing the nitrate assimilation gene cluster. We found that r/m values (the ratio of nucleotide changes due to recombination relative to point mutation) were well in excess of 1 for both the nitrate assimilation genes and the core genomic regions that flank them (Table 1), indicating that homologous recombination has a role in modulating diversity within these genomic regions. It is being increasingly acknowledged that bacteria can behave like sexually recombining populations (Fraser et al. 2007) and that recombination has a role in maintaining overall population diversity and in facilitating gene-specific, rather than genome-wide, sweeps within populations (Shapiro et al. 2012; Shapiro 2016; Rosen et al. 2015). High rates of homologous recombination are likely a general feature of free-living marine bacterial populations (Vergin et al. 2007; Vos and Didelot 2009), including *Prochlorococcus*.

**Table 1.** Rates of recombination relative to mutation for representative genomic regions in high-light and low-light adapted *Prochlorococcus*.

| Region | Functional Group | Sequences in Alignment | Alignment Length (bp) | $\kappa$ | $\delta$ | $\nu$ | R/$\theta$ | $\rho/\theta$ | r/m |
| --- | --- | --- | --- | --- | --- | --- | --- | --- | --- |
| **High-light adapted *Prochlorococcus* (HLII and HLVI clades)** |  |  |  |  |  |  |  |  |  |
| *nirA-moaA* | nitrate | 33 | 11428 | 3.32 | 4202 | 0.012 | 0.68 | 1.36 | 34.0 |
| *polA* region<sup>a</sup> | core flanking | 19 | 11554 | 3.72 | 8035 | 0.020 | 0.20 | 0.41 | 33.2 |
| **Low-light adapted *Prochlorococcus* (LLI clade)** |  |  |  |  |  |  |  |  |  |
| *moaB-narB* | nitrate | 17 | 11985 | 2.91 | 435 | 0.021 | 1.78 | 3.57 | 16.5 |
| *pyrG* region<sup>b</sup> | core flanking | 15 | 10080 | 2.81 | 964 | 0.029 | 2.17 | 4.34 | 59.7 |
| *ppk* region<sup>c</sup> | core flanking | 15 | 11918 | 3.19 | 1056 | 0.031 | 0.93 | 1.85 | 30.3 |

a 3' flanking region, containing *polA*, adjacent to the nitrate assimilation gene cluster in the HLII clade.
b 5' flanking region, containing *pyrG*, adjacent to the nitrate assimilation gene cluster in the LLI clade.
c 3' flanking region, containing *ppk*, adjacent to the nitrate assimilation gene cluster in the LLI clade.
$\kappa$ Transition/transversion rate ratio as estimated by PhyML under the HKY85 model.
$\delta$ Mean length of DNA imported by homologous recombination.
$\nu$ Divergence rate, per site, of DNA imported by homologous recombination.
R/ $\theta$ Per-site rate of initiation of recombination relative to the population mutation rate.
$\rho/\theta$ Population recombination rate relative to the population mutation rate ( $\rho=2R$ ).
r/m Relative impact of recombination versus mutation on the per-site substitution rate. Equal to $(R/\theta) \times \delta \times \nu$ .

To further evaluate these processes in *Prochlorococcus*, we next assessed the cross-habitat diversity of the nitrate assimilation genes in comparison to the core gene, *gyrB*, and the phosphate assimilation genes, *pstB* and *pstS*. The latter are examples of genes that are known to experience high rates of recombination, horizontal gene transfer, and/or efficient selection in *Prochlorococcus* populations (Coleman and Chisholm 2010). Comparing single cells from two populations (North Pacific vs. North Atlantic), we found no significant phylogenetic divergence in the core gene *gyrB* and in genes in the upstream half of the nitrate assimilation pathway (Table 2 and Fig. 5). As expected, the *pstB* and *pstS* genes did cluster phylogenetically based on the populations from which they were derived (Table 2 and Fig. 5). Notably, the nitrite reductase gene *nirA* exhibited significant phylogenetic divergence between the two populations in 2 out of 3 subsampled data sets (Table 2). This gene may have experienced high enough recombination rates to facilitate its phylogenetic divergence between geographically distant populations or may have been fine-tuned by selection for optimal function in these contrasting ecosystems.

**Fig. 5.**
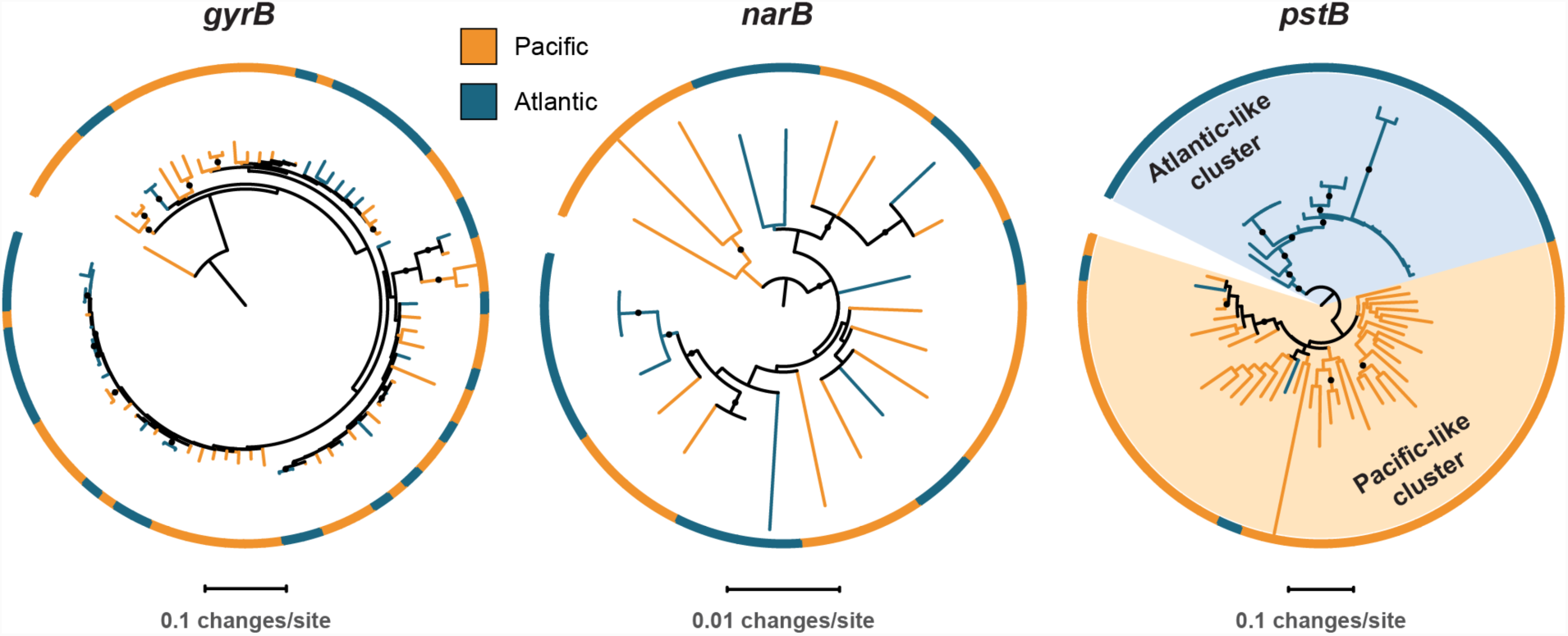
Representative phylogenetic patterns for DNA gyrase subunit B (*gyrB*) in comparison to nitrate reductase (*narB*) and the phosphate assimilation gene *pstB* (encoding the ABC transporter ATP binding subunit) from single cells in surface populations at HOT (Hawai’i Ocean Time-series) and BATS (Bermuda Atlantic Time-series Study). The *gyrB* and *narB* genes do not exhibit significant phylogenetic divergence between the two sites. The *pstB* gene, in contrast, has significantly diverged into Atlantic-like and Pacific-like clusters of sequences due to frequent recombination, gene transfer, and/or selection (sensu Coleman et al. 2010).

**Table 2.**
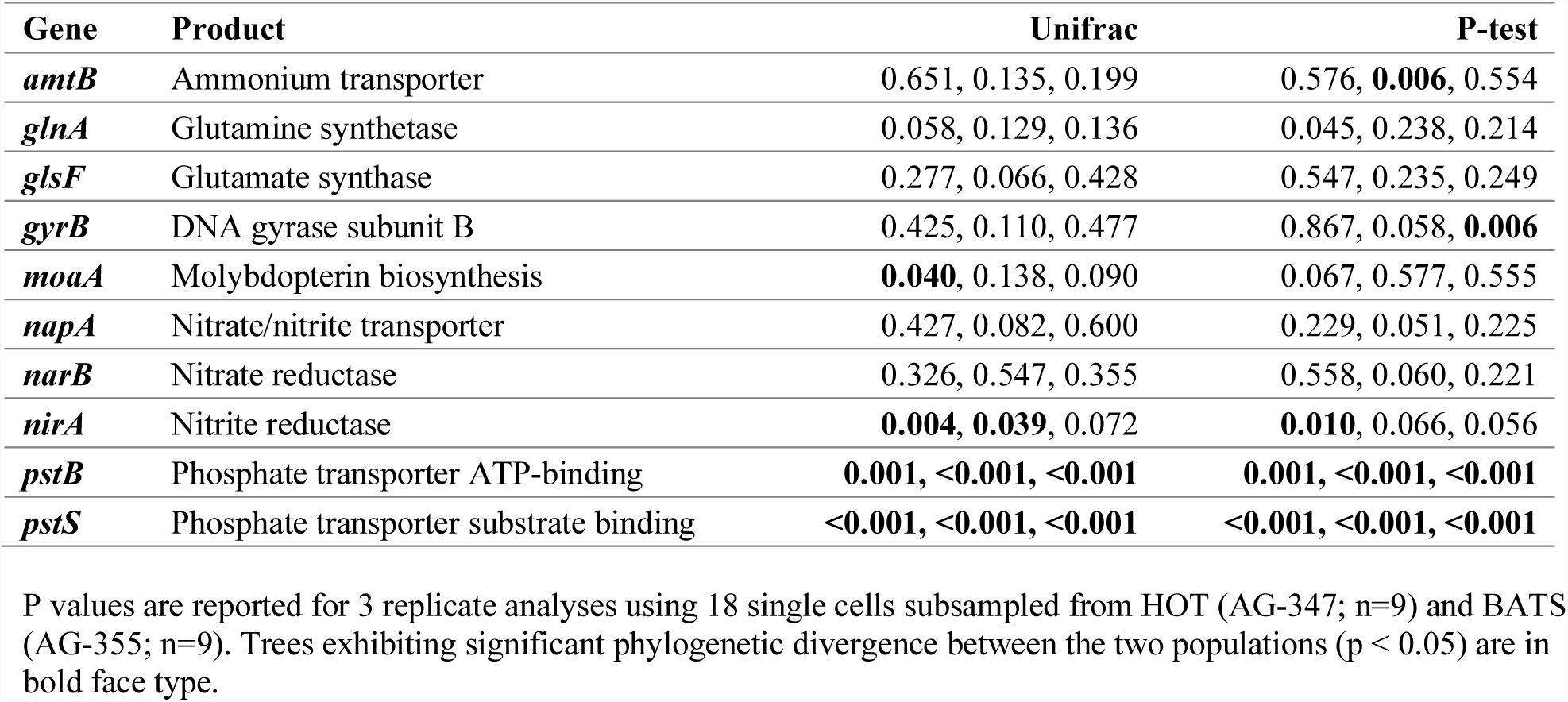
Significance (p value) for divergent phylogenetic gene clusters separating populations in the North Pacific Subtropical Gyre (HOT) and in the North Atlantic Subtropical Gyre (BATS).

While it generally appears that recombination and selection have not been sufficient to drive significant phylogenetic divergence between populations for most nitrate assimilation genes (Table 2 and Fig. 5), we did observe a particularly high nucleotide similarity between these genes within clades (Fig. 6). Homologous recombination within the nitrate assimilation gene clusters of two closely related genomes can serve as a cohesive force that reinforces genetic similarity. Further, tests of adaptive evolution indicated that the nitrate assimilation genes are subject to strong purifying selection (Supplementary Tables 1-3) which would act to constrain divergence at non-synonymous sites. Homologous recombination between existing nitrate assimilation gene clusters is expected to have a complementary role in maintaining the observed phylogenetic cohesion within individual clades of *Prochlorococcus* (Fig. 6).

**Fig. 6.**
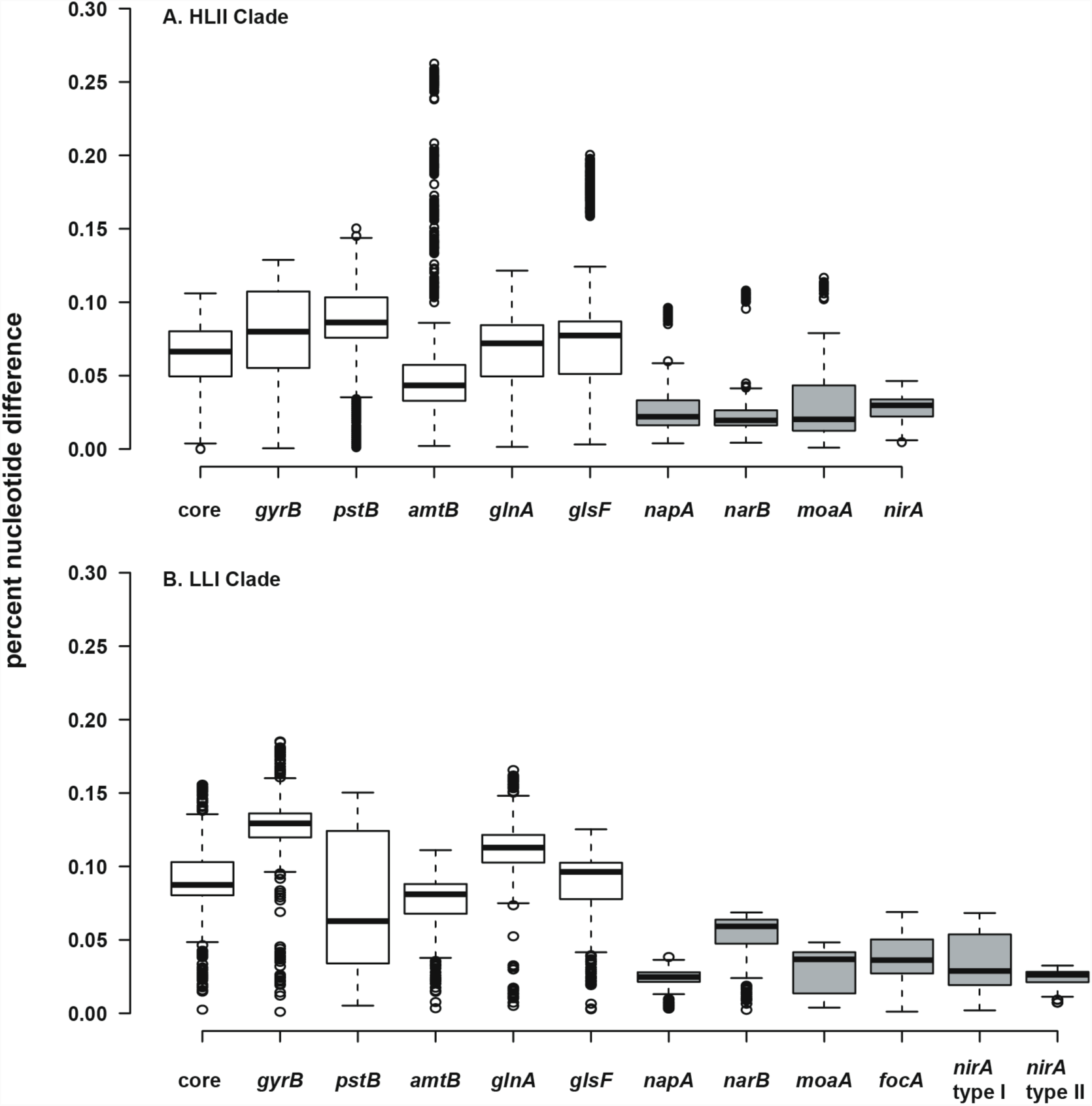
Percent nucleotide difference of selected genes for the HLII (A) and LLI (B) clades of *Prochlorococcus*. The ‘core’ genes are a concatenated alignment of up to 37 PhyloSift marker genes. Genes in the nitrate assimilation cluster are in gray. Center lines show the medians, box limits indicate the 25th and 75th percentiles, whiskers extend 1.5 times the interquartile range from the 25th and 75th percentiles, and outliers are represented by dots. For HLII genes (A), n = 5151, 4186, 3321, 3570, 4371, 3240, 378, 351, 351, 351 sample points. For LLI genes (B), n = 496, 406, 435, 435, 406, 406, 153, 136, 136, 171, 210, 28 sample points.

Homologous recombination in shared genomic regions that flank the nitrate assimilation gene cluster has likely facilitated both the loss and acquisition of these genes within clades. Given that rates of loss are expected to be several orders of magnitude faster than those of gains (Apagyi et al. 2018), the patchy distribution of the nitrate assimilation trait across the phylogeny of individual *Prochlorococcus* clades is more likely dominated by gene loss. Low nucleotide diversity among nitrate assimilation genes within clades (Fig. 6), also suggests that replacement and/or acquisition of alleles through homologous recombination has driven occasional sweeps of nitrate assimilation alleles through populations of closely related *Prochlorococcus*. Overall, we expect that the maintenance of this trait in wild populations is a consequence of the higher relative fitness of nitrate assimilating cells under conditions of overall nitrogen limitation (Berube et al. 2016). It is also likely that frequency-dependent selection processes (Cordero and Polz 2014) have played a role in setting the equilibrium between genotypes that possess the nitrate assimilation genes and those that lack them.

### Macroevolutionary implications of differential trait variability in *Prochlorococcus* clades and ecotypes

Our gene content, phylogenomic, and diversity analyses suggest that the evolution of nitrate assimilation in *Prochlorococcus* largely proceeds through vertical descent, stochastic gene loss, and homologous recombination between closely related cells. However, this leaves several patterns unexplained. In the absence of horizontal gene transfer between distant taxa, how can a trait, that is present in recently branching lineages, be absent in more deeply branching lineages? We addressed this by looking to the broader physiological and ecological context of *Prochlorococcus’* evolution. As discussed above, selection appears to favor nitrate assimilation in conditions where the supply of light is enhanced and the overall supply of nitrogen is low (Berube et al. 2016). Since the trait appears to have descended vertically in *Prochlorococcus* as a whole, this raises the possibility that basal LL clades, which lack nitrate assimilation and dominate at depth in the contemporary oceans, are derived from ancestral populations that had a broader depth distribution. This is consistent with the argument that the long-term evolution of *Prochlorococcus* proceeded via a sequence of niche-constructing, adaptive radiations (Braakman et al. 2017). In this model of *Prochlorococcus’* evolution, metabolic innovations enhanced overall nutrient affinity by increasing the photosynthetic electron flux, thereby facilitating nutrient uptake at ever lower nutrient levels and making new realized niches available to *Prochlorococcus* (Braakman et al. 2017). It is inferred that each new ecotype increased the harvesting of solar energy in the brightly illuminated but nutrient poor surface waters, thereby serially drawing down nutrient levels and restricting basal lineages to deeper waters (Braakman et al. 2017).

In our proposed macroevolutionary scenario, nitrate assimilation genes descend vertically from ancestral populations and stochastic loss of these genes is balanced by the fitness benefit of nitrate assimilation in environments where it offers a selective advantage. Niche differentiation between *Prochlorococcus* ecotypes could result in basal lineages losing nitrate assimilation genes over time as they became specialized to niches characterized by adaptation to lower irradiances (Fig. 7). In the basal LLIV clade, which is restricted to the deepest region of the euphotic zone, loss of these genes appears to have run to completion rather than reaching some frequency-dependent equilibrium that is observed in the recently emerged clades. Homologous recombination would be limited to closely related cells which share sufficient nucleotide similarity to facilitate appreciable rates of replacement and/or acquisition of nitrate assimilation alleles (Fig. 7).

**Fig. 7.**
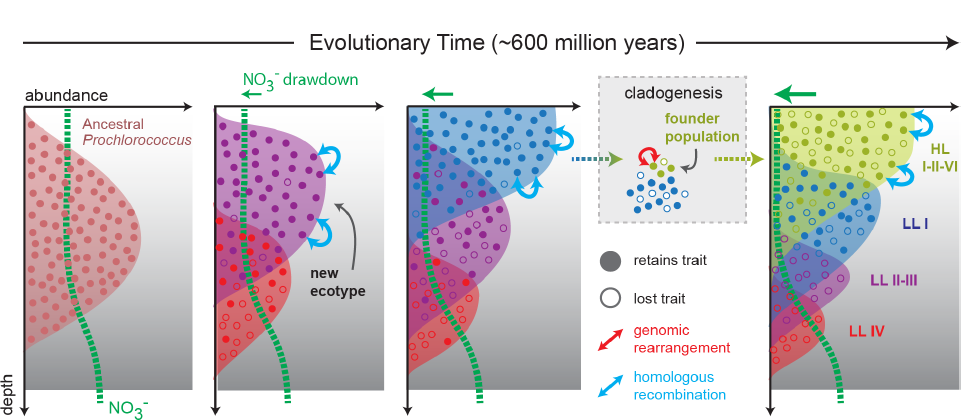
Proposed model mapping the vertical inheritance of the nitrate assimilation pathway onto a general model of speciation in *Prochlorococcus* (Braakman et al. 2017). We assume different cost:benefit ratios at top and bottom of the water column based on the energetic requirements of the nitrate assimilation pathway and the selective advantage of carrying this trait in environments with low nitrogen availability (see main text for further discussion). We further assume that ancestral *Prochlorococcus*, similar to *Synechococcus*, were capable of nitrate assimilation. As new *Prochlorococcus* clades/ecotypes emerged to more efficiently harness light energy and facilitate the draw-down of nitrogen at the surface, basal lineages were partitioned to the deeper regions of the euphotic zone (Braakman et al. 2017). In these basal lineages, higher relative costs of the pathway combined with access to other nitrogen sources (e.g. amino acids) has hastened the loss of this trait through stochastic gene loss. In more recently emerging lineages (LLI, HLI, HLII, HLVI), the trait has been retained with intra-clade frequencies influenced by the specific chemical and physical characteristics of the environment in which they are found (Berube et al. 2016). The founder effect drives punctuated changes (e.g. genome-wide rearrangements) during speciation, while homologous recombination acts to constrain the divergence of gene sequence and order within clades.

While we cannot discount the possibility of non-homologous based gene acquisition in different genomic regions during speciation, the genomic rearrangement of nitrate assimilation genes in the most recently emerging lineages (Fig. 4) is consistent with brief periods of enhanced genetic drift due to the founder effect (Hallatschek et al. 2007). Genomes within individual *Prochlorococcus* clades are highly syntenic, but large-scale genomic rearrangements exist between the genomes of different clades (Yan et al. 2018). Ecological differentiation of *Prochlorococcus* likely involved a period of rapid population expansion as new ecotypes gained access to a previously inaccessible nutrient pool (Braakman et al. 2017). Given the size of *Prochlorococcus* populations, any bottleneck at the front of this expansion is not expected to have been severe, but it has been observed that large genomic rearrangements can become fixedin bacterial populations even under weak bottlenecks and in the absence of horizontal gene transfer (Raeside et al. 2014). The increased likelihood of genome-wide genomic rearrangements following speciation events is a plausible explanation for the existence of nitrate assimilation gene clusters in different genomic locations when comparing between clades.

### Conclusions

We argue that the diversity and intra-specific distribution of the nitrate assimilation trait in *Prochlorococcus* is likely driven by a combination of vertical inheritance and gene loss, and rarely due to horizontal gene acquisition driven by non-homologous recombination. These processes have largely resulted in a patchy distribution of this trait among cells within clades that harbor nitrate assimilation genes. Homologous recombination further acts to constrain the divergence of this trait within clades and may have promoted gene-specific selective sweeps within populations of closely related cells. Seasonality in environmental conditions, under which this trait likely confers a selective advantage to cells (Berube et al. 2016), has probably facilitated the maintenance of this patchy trait distribution. We further propose that speciation and niche partitioning involved brief periods of enhanced drift and rapid change that facilitated the genomic rearrangement of nitrate assimilation genes. The restriction of basal *Prochlorococcus* lineages to greater depths during the general process of niche partitioning explains their loss of the nitrate assimilation trait (Fig. 7). Overall, the underlying dynamics governing the loss and retention of nitrate assimilation within clades and during the genesis of new clades are closely intertwined. Superficially, this emergent pattern in microbial trait variability might appear to be the result of horizontal gene transfer, but our evidence indicates that the observed patterns can be attributed to processes of vertical descent, gene loss, and recombination between close relatives that have operated throughout the entire radiation of *Prochlorococcus*.

## METHODS

### Dataset

A set of 463 *Prochlorococcus* and *Synechococcus* single cell genome assemblies (Berube et al. 2018) were filtered to exclude 76 genomes with less than 25% genome recovery, as determined by checkM (Parks et al. 2015). The AG-363-L17, AG-418-C09, andAG-418-C17 single cell genome assemblies were excluded because they contained both bacterial and phage genomes. A subset of 206 of the remaining single cells were screened using a PCR assay (Supplementary Methods) to confirm the presence or absence of *narB* in the amplified single cell DNA. An additional 33 single cell genome assemblies had annotated *narB* genes, but were not screened by PCR. This resulted in a total of 239 single cells with empirical evidence for the presence or absence of *narB*, and were retained in the data set; these single cell genome assemblies had a median genome recovery of 79%. The final data set encompassed 321 genomes after inclusion of reference culture genomes for 61 *Prochlorococcus* (26 closed and 35 permanent draft) and 21 *Synechococcus* (14 closed and 7 permanent draft). Eight single cells, belonging to the HLIII and HLIV clades of *Prochlorococcus* (Malmstrom et al. 2013), were additionally used as references for the core marker gene phylogeny only. All genomes were downloaded from the Integrated Microbial Genomes (IMG) system (Chen et al. 2017). ProPortal CyCOGs v6.0 definitions (Berube et al. 2018) were used as primary annotations. IMG accession numbers for the 321 genomes in the final data set are provided in Supplementary Table 4.

### Phylogenetic inference

For core gene/protein phylogenies, we used PhyloSift (Darling et al. 2014) to search the genome assemblies in our data set for 37 marker gene families (Darling et al. 2014) and generate concatenated nucleotide codon alignments for these marker genes. The alignments were manually curated to remove poorly aligned regions and translated to generate amino acid alignments. Maximum likelihood trees were then generated using RAxML 8.2.9 (Stamatakis 2014) with automatic bootstopping criteria enabled and using the following command line parameters: raxmlHPC-PTHREADS-AVX -T 20 -f a -N autoMRE -m PROTGAMMAAUTO (amino acid alignments); raxmlHPC-PTHREADS-AVX -T 20 -f a -N autoMRE -m GTRCAT (nucleotide alignments).

For individual genes belonging to specific CyCOG families, we used MACSE (Ranwez et al. 2011) to generate nucleotide codon alignments. The alignments were manually curated to remove taxa with >80% missing data. For taxa with genes breaking across contig boundaries, the longest aligning segment was retained. The nucleotide alignments were translated to generate amino acid alignments and maximum likelihood trees were generated with RAxML 8.2.9 (Stamatakis 2014), using and the command line parameters described for core protein marker phylogenies. Percent nucleotide difference for genes belonging to HLII and LLI taxa were determined using MOTHUR (Schloss et al. 2009). Genes belonging to these clades were extracted from the codon alignments and the dist.seqs command in MOTHUR (Schloss et al.2009) was used to create a column distance matrix for each gene. BoxPlotR (Spitzer et al. 2014) was used for visualization.

### Genome location and synteny

Single cell genome assemblies were filtered for contigs greater than 40 kbp in length that contained complete nitrite and/or nitrate assimilation gene clusters. This yielded contigs with sequence data extending from the nitrate assimilation gene cluster into adjacent genomic regions for 1 HLI, 26 HLII, 2 HLVI, and 18 LLI single cell genomes. Two HLII and 10 LLI culture genomes with nitrite and/or nitrate assimilation gene clusters were also included. Contigs were then aligned to closed reference genomes using the progressiveMauve algorithm (Darling et al. 2010) in MAUVE 2.4.0 with default parameters.

Alignments for contigs derived from high-light adapted single cells used the following reference genomes: MIT9301 (HLII), AS9601 (HLII), MED4 (HLI), and MIT9515 (HLI). Alignments for contigs derived from low-light adapted single cells used the following reference genomes for the LLI clade: NATL2A, MIT0915, and MIT0917.

### Analysis of covariation in gene content

HLII and LLI single cells were filtered to exclude those with < 75% genome recovery, as determined by checkM (Parks et al. 2015), yielding 105 single cells (83 HLII and 22 LLI single cells). Each group of genomes had median genome recoveries of 90% and 87%, respectively. The number of each CyCOG in each genome was enumerated and both abundance and binary (presence/absence) matrices were created for each set of HLII and LLI genomes. CyCOGs either shared by all genomes or exhibiting low representation among genomes have little information content with regards to the identification of co-varying genes. Thus, core CyCOGs and CyCOGs found in fewer than 3 genomes were excluded from analysis. Core CyCOGs in partial single cell genome assemblies were operationally defined as CyCOGs found in at least M% of genomes, where M% is the median genome recovery for each set of genomes (for example, CyCOGs found in at least 19 (22 x 0.87) LLI genomes were estimated to be core CyCOGs). Binary matrices were imported into MORPHEUS (Broad Institute; MA, USA, accessed at https://software.broadinstitute.org/morpheus/) and CyCOGs were hierarchically clustered based on the Jaccard distance measure and using average linkage to identify CyCOGs that co-vary with the nitrate assimilation gene cluster. Gene enrichment analysis was also used to examine covariation of CyCOGs with the nitrate assimilation genes. Over- and under-representation of CyCOGs were evaluated using BiNGO 3.0.3 (Maere et al. 2005) in Cytoscape 3.4 (Shannon et al. 2003). For each set of 83 HLII and 22 LLI single cells, genes found in genomes containing *narB* were evaluated against the set of genes found in all genomes in the set. Significant enrichment of flexible genes in the test set was assessed using the hypergeometric statistical test and the Benjamini and Hochberg correction for multiple hypothesis testing.

### Recombination detection

Complete nitrate assimilation gene clusters and core flanking regions adjacent to the nitrate assimilation gene clusters were extracted from Mauve alignments. Nucleotide alignments for these genomic tracts were manually curated and PhyML 3.0 (Guindon et al. 2010) was used to construct a starting tree and to estimate the transition/transversion ratio (kappa) using default parameters. The impact of recombination relative to mutation was then assessed with CLONALFRAMEML (Didelot and Wilson 2015) using the nucleotide alignment, phylogenetic tree, and estimated kappa value as inputs. The ρ*/*θ value is a measure of the occurrence of initiation or termination of recombination relative to the population mutation rate. The relative impact of recombination versus mutation (r/m) is measured by accounting for the length (δ) and genetic distance (ν) of the recombining fragments (Didelot and Wilson 2015).

### Beta diversity analysis

Gene sequences belonging to HLII clade single cells derived from two surface populations (AG-347, HOT, Hawai’i Ocean Time-series, 5m depth; and AG-355, BATS, Bermuda Atlantic Time-series Study, 10m depth) were extracted from gene alignments. These two environmental samples had the highest sample number of HLII clade single cells and included a minimum of 9 sequence representatives for each population. Nine sequences were subsampled without replacement from each population using MOTHUR (Schloss et al. 2009) to yield a total of 18 sequences in the final alignment. The subsampled alignment was used to build a phylogenetic tree using RAxML 8.2.9 (Stamatakis 2014) with the GTRCAT model and automatic bootstopping criteria enabled. Trees were rooted at the midpoint using PyCogent (Knight et al. 2007). Beta diversity was assessed using Fast Unifrac (Hamady et al. 2010) as implemented in PyCogent (Knight et al. 2007) to determine significance values based on Unifrac and the P-test (Martin 2002): fast_unifrac_permutations_file(tree_in, envs_in, weighted=False, num_iters=1000, test_on=“Pairwise”); fast_p_test_file(tree_in, envs_in, num_iters=1000, test_on=“Pairwise”). This analysis was repeated for 3 independent subsampled data sets.

## Supporting information

## ACKNOWLEDGEMENTS

This work was supported by grants from the National Science Foundation (OCE-1153588 and DBI-0424599 to S.W.C. and OCE-1335810 to R.S.), the Simons Foundation (Life Sciences Project Award IDs 337262 and 509034SCFY17, S.W.C; SCOPE Award ID 329108, S.W.C), and the Gordon and Betty Moore Foundation (Grant IDs GBMF495 and GBMF4511 to S.W.C.). This paper is a contribution from the Simons Collaboration on Ocean Processes and Ecology (SCOPE) and the NSF Center for Microbial Oceanography: Research and Education (C-MORE).

## AUTHOR CONTRIBUTIONS

P.M.B. conceived and designed the study; P.M.B. and A.R. designed and performed the *narB* PCR screen; P.M.B. and R.S. analyzed data; R.B. and P.M.B. developed the macroevolution model; P.M.B., A.R., R.B., R.S., and S.W.C. wrote the paper.

## COMPETING INTERESTS

The authors declare no competing financial interests.

## ADDITIONAL INFORMATION

Supplementary information accompanies this paper.

