## Supplementary material for "Emergence of trait variability through the lens of nitrogen assimilation in *Prochlorococcus*"

#### Supplementary Methods

**Genome sequencing.** Genomes for the following cultured LLI *Prochlorococcus* strains were sequenced as part of this study: MIT0912, MIT0913, MIT0915, and MIT0917. The MIT0912 and MIT0913 strains were isolated in July 2009 from Station ALOHA (Hawai'i Ocean Time-series, 23.75$^o$N, -158$^o$E) from a depth of 175m on the HOT212 cruise (KM0915). MIT0915 and MIT0917 were derived from the P0902-H212 and P0903-H212 enrichment cultures, respectively (Berube et al. 2016). Cells were grown to mid exponential phase and pelleted by centrifugation. DNA was isolated by phenol/chloroform extraction (Wilson 2001). PacBio library preparation and sequencing was carried out by the MIT BioMicro Center and the UMass Worcester Medical School's Deep Sequencing Core Facility. Assembly of PacBio reads was performed using the hierarchical genome-assembly process (Protocol = RS_HGAP_Assembly.2) as implemented in SMRT Analysis 2.3.0 (Chin et al. 2013) with the following parameters adjusted: Minimum Polymerase Read Quality = 0.85 and Genome Size = 2000000 bp (default settings were used for all other parameters). Overlapping ends of the assembly were identified using BLAST and the assembly was manually circularized. Circular assemblies were corrected using the RS_Resequencing.1 protocol in SMRT Analysis 2.3.0 (Chin et al. 2013) with the following parameters: Minimum Polymerase Read Quality = 0.85 and Consensus Algorithm = Quiver. These genomes were deposited with IMG (accession numbers: 2681812899, 2681812900, 2681812901, 2681812859), annotated using IMG Annotation Pipeline version 4 (Markowitz et al. 2014; Chen et al. 2017), and included in ProPortal CyCOGs 6.0 (Berube et al. 2018).

**Screening assay for *narB*.** We used a PCR assay to screen for the presence/absence of the *narB* gene in the amplified DNA from sorted single cells. The design of the PCR primers was based on an alignment of 49 *narB* sequences extracted from 13 marine *Synechococcus* genomes

(WH7805, WH7803, BL107, CC9902, CB0205, WH8102, CC9605, CB0101, RS9916, WH8016, CC9311, RCC307, and WH8109), 1 *Prochlorococcus* contig derived from an environmental metagenome assembly (Astorga-Eló et al. 2015), 11 LLI *Prochlorococcus* genomes (PAC1, MIT0915, MIT0917, AG-363-M20, AG-402-L20, AG-402-M23, AG-402-O21, AG-311-K16, AG-402-L09, AG-341-O20, and AG-331-D10), 1 HLII *Prochlorococcus* genome (AG-335-O19), 22 HLII *Prochlorococcus* genomes (SB, MIT0604, AG-355-I20, AG-402-I23, AG-347-I22, AG-335-I15, AG-347-G18, AG-355-G23, AG-347-M23, AG-347-O22, AG-347-I19, AG-347-I21, AG-355-N18, AG-402-K16, AG-347-G20, AG-347-K17, AG-347-L20, AG-355-B23, AG-355-J09, AG-402-L23, AG-402-G23, and AG-402-K22), and 1 HLVI *Prochlorococcus* genome (AG-363-B18). Sequences were aligned by codon using MACSE (Ranwez et al. 2011) and the following primers were selected: narB.SAG.Deg.1786F (5'-CANTGGCAYACNATGAC-3') and narB.SAG.Deg.2004R (5'-RAANCCCCARTGCATNGG-3'). The forward primer has a 32-fold degeneracy targeting a conserved region encoding the [Q/H]WHTMT polypeptide, and the reverse primer has a 64-fold degeneracy targeting a conserved region encoding the PMHWGF polypeptide. These motifs are conserved between the *Prochlorococcus* and marine *Synechococcus* genomes in our data set and are expected to amplify most *narB* sequences within these groups. PCR conditions were optimized using *Prochlorococcus* strains SB and MIT0917. The reaction consisted of 1 µM forward primer, 1 µM reverse primer, 2 µl of template DNA, 200 µM dNTPs, 1x Phusion HF Buffer (New England BioLabs), 1 unit Phusion High-Fidelity DNA Polymerase (New England BioLabs), and 0.2x SYBR Green (Lonza) in a 30 µl total volume. Real-time PCR was performed using a Bio-Rad CFX96 instrument programmed for 40 cycles of 98°C for 10s, 54°C for 30s, and 72°C for 30s. Positive amplification was identified by the presence of a single peak in the melting curve between 78-83°C and by visual inspection for an amplicon length of 219 bp on an agarose gel.

While PCR can identify genomic regions with poor sequencing coverage, biases in DNA amplification can result in gene copy numbers that still fall below the detection limit of PCR. Thus, we assessed false negative rates using a set of 4 reference cultures: *Prochlorococcus* SB (HLII clade, *narB*$^+$), *Prochlorococcus* MIT9301 (HLII clade, *narB*$^-$), *Prochlorococcus* MIT0917 (LLI clade, *narB*$^+$), and *Prochlorococcus* NATL2A (LLI clade, *narB*$^-$). Each culture was grown to mid-exponential phase, at which point, cell concentrations were determined by flow cytometry using a GUAVA flow cytometer (Millipore). The cultures were combined at equivalent concentrations, preserved using 10% glycerol, flash frozen in liquid nitrogen, and stored at -80°C. Single cells were sorted and the DNA was amplified using the WGA-X method (Stepanauskas et al. 2017) by the Bigelow Laboratory Single Cell Genomics Center. ITS sequences were amplified and sequenced using the Sanger method to associate wells on the 384 well plate with each control strain. The *narB* PCR assay was executed on the amplified single cell DNA from the control strains. Template DNA was prepared by making a 50-fold dilution of a subsample of the amplified DNA in 1x TE buffer. The false negative rate (likely due to extremely poor genome amplification across the nitrate assimilation gene cluster) was 8% for the control strains containing *narB*. We did not observe any false positive results for control strains lacking *narB*.

Our *narB* PCR assay was then applied to 6 experimental plates of amplified single cell DNA: AG-347, AG-355, AG-363, AG-402, AG-418, and AG-459. A SAG was deemed positive if PCR resulted in a narrow peak in the melting curve at approximately 78-83$^{o}$C. Those with no amplification or with a melting curve characteristic of primer dimers were deemed negative. Twelve amplicons that were either borderline or potentially spurious were sequenced using the Sanger method to confirm whether or not they were *narB*. Finally, the results of the PCR screen were used to select an additional 47 *narB*$^{+}$ SAGs for whole genome sequencing, annotation by IMG (Markowitz et al. 2014), and inclusion in ProPortal CyCOGs v6.0 (Berube et al. 2018).

Comparison of PCR screening results with CyCOG annotations for the partial single cell genome assemblies supports the effectiveness of PCR as a method to assess the presence/absence of functional genes in large single cell genomics data sets. Of the 206 single cells screened by PCR, the PCR data and the annotation data agreed in 88% of cases. For single cells with an annotated *narB* gene (61/206 cells), 97% tested positive for the presence of *narB* by PCR (59/61 cells). For single cells that tested positive for the presence *narB* by PCR (81/206), 73% had an annotated *narB* gene (59/81 cells). The median genome recovery of these 81 single cell genomes was 80%, providing an expected occurrence of 65 single cell genomes that should contain an annotated *narB* gene. A post hoc analysis comparing 59 observed to 65 expected occurrences of genomes containing an annotated *narB* gene suggests an 9% false negative rate for the PCR screening assay relative to the 8% false negative rate determined using control strains.

**Estimates of dN/dS and tests of adaptive evolution.** The CODEML application of the PAML package (Yang 1997; Yang 2007) was used to estimate synonymous and nonsynonymous substitution rates (dN/dS) and assess the likelihood of adaptive evolution for a selection of protein coding genes (*gyrB; pstB; amtB; glnA; glsf; napA; narB; moaA; focA; nirA*). The CODEML parameters employed were as described by Jeffares et al. (Jeffares et al. 2015) except for fixing branch lengths (fix_blength=2) and using a small difference value (Small_Diff) of 1e-7 in site model tests and 1e-8 in branch-site model tests. Initial data sets were the codon alignments used for the gene and protein phylogenies described in the main text. Given the large number of taxa in the codon alignments and the high computational demands of CODEML, we found it necessary to identify representative taxa using MOTHUR (Schloss et al. 2009) by clustering the sequences with a distance cutoff of 0.01 using the following commands:

```
mothur unique.seqs(fasta=alignedcodons.fasta)

mothur dist.seqs(fasta= alignedcodons.unique.fasta)

mothur cluster(column= alignedcodons.unique.dist, name=
alignedcodons.names, method=opti, cutoff=0.01)

mothur get.oturep(column= alignedcodons.unique.dist, name=
alignedcodons.names, list= alignedcodons.unique.opti_mcc.list)
```

The representative taxa identified by MOTHUR were extracted from the original codon alignments and RAxML (Stamatakis 2014) was used to infer the phylogeny with the following parameters: `raxmlHPC-PTHREADS-AVX -T 20 -f a -x -p -N 100 -m GTRCAT`

Site model tests used the following CODEML control file:

```
      seqfile = <codon alignment for gene>
     treefile = <RAxML besttree file>
      outfile = outfile.txt
        noisy = 9
      verbose = 1
      runmode = 0
      seqtype = 1
    CodonFreq = 2
        ndata = 1
        clock = 0
        model = 0
      NSsites = 0 1 2 7 8
        icode = 0
    fix_omega = 0
        omega = .4
   Small_Diff = 1e-7
    cleandata = 0
  fix_blength = 2
```

dN/dS (omega) values and log-likelihoods for each of the 5 models were extracted from the output file. Significance was assessed using the likelihood ratio test statistic and the chi-squared distribution.

Branch-site model tests used the following CODEML control file for the null hypothesis of adaptive evolution across sites and lineages (Hypothesis H0). The branch leading to the lineage under examination (foreground branch) was marked with '$1' in the RAxML bestree file:

```
    seqfile = <codon alignment for gene>
   treefile = <Marked RAxML besttree file>
    outfile = outfile.txt
      noisy = 9
    verbose = 1
    runmode = 0
    seqtype = 1
  CodonFreq = 2
      ndata = 1
      clock = 0
      model = 2
    NSsites = 2
      icode = 0
  fix_omega = 1
      omega = 1
 Small_Diff = 1e-8
  cleandata = 0
fix_blength = 2
```

The hypothesis for positive selection among some sites in the foreground branch (Hypothesis H1) was then examined by using the same control file but with <`fix_omega = 0`> to allow for a class of sites in the foreground branch with dN/dS > 1.

dN/dS values for each site class in the foreground and background branches as well as log-likelihoods for the two hypotheses (H0 vs. H1) were extracted from their respective output files. Significance was assessed using the likelihood ratio test statistic and the chi-squared distribution. Although the likelihood ratio test does not follow the chi-squared distribution in the case of branch-site tests of adaptive evolution, the chi-squared critical value for 1 degree of freedom was used to make the test conservative (Jeffares et al. 2015).

### Supplementary Information Figures

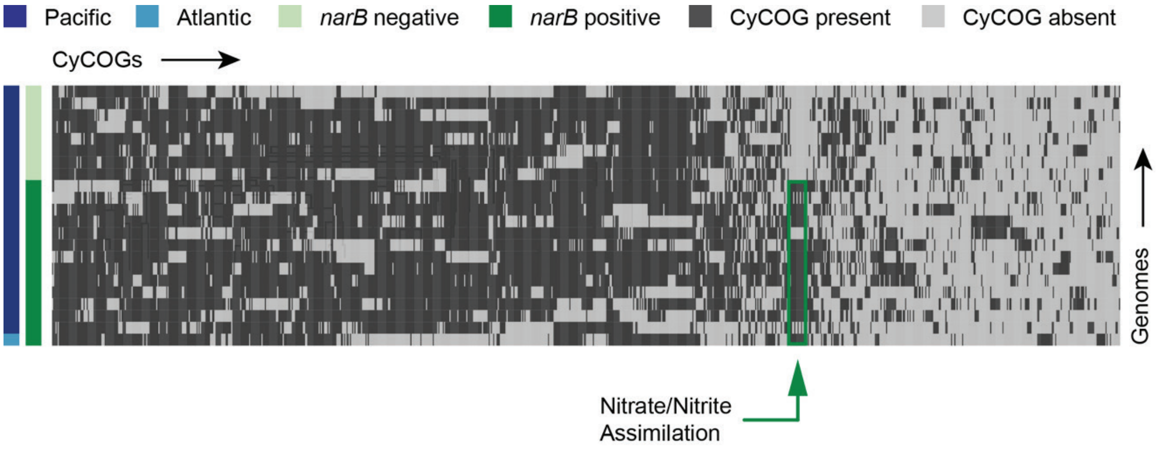


**Supplementary Fig. 1.** Hierarchical clustering of presence and absence distributions for flexible CyCOGs found in 22 *Prochlorococcus* single cell genomes belonging to the LLI clade with genome recoveries of at least 75% (median 87%). Genomes are sorted by Atlantic and Pacific Oceans and by the presence/absence of the *narB* gene – a marker for the capacity to assimilate nitrate. Other than genes in the nitrate assimilation gene cluster (green box), no CyCOGs were over- or under-represented among the flexible genes in genomes containing *narB*.

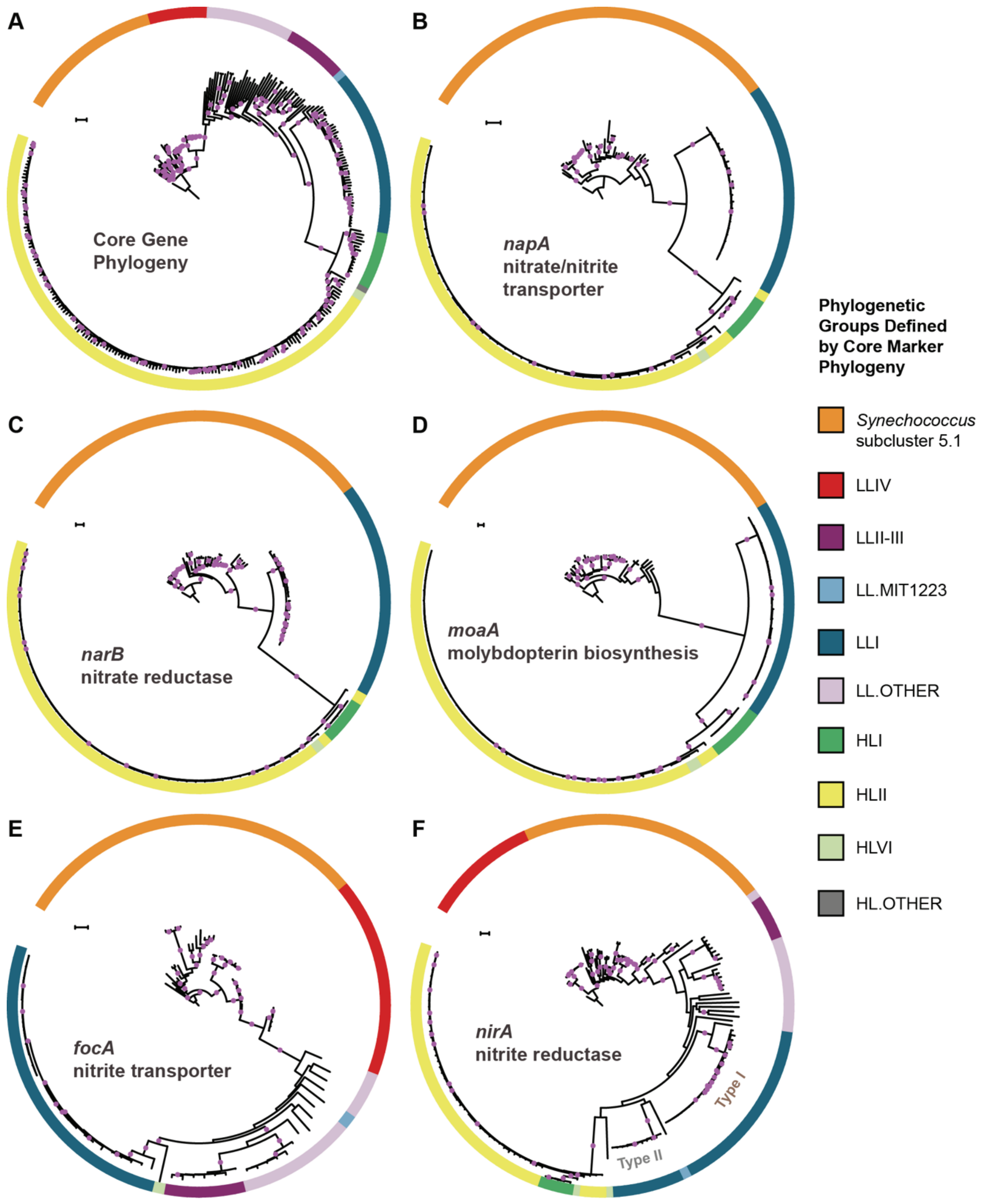


**Supplementary Fig. 2.** The core marker gene phylogeny of *Prochlorococcus* and *Synechococcus* (A) in comparison to gene phylogenies for the nitrate/nitrite transporter, *napA* (B), the nitrate reductase, *narB* (C), the molybdopterin biosynthesis protein, *moaA* (D), the nitrite transporter, *focA* (E), and the nitrite reductase, *nirA* (F). Filled purple circles on branches indicate that the associated taxa clustered together in at least 75% of trees. Scale bars are 0.1 changes per site.

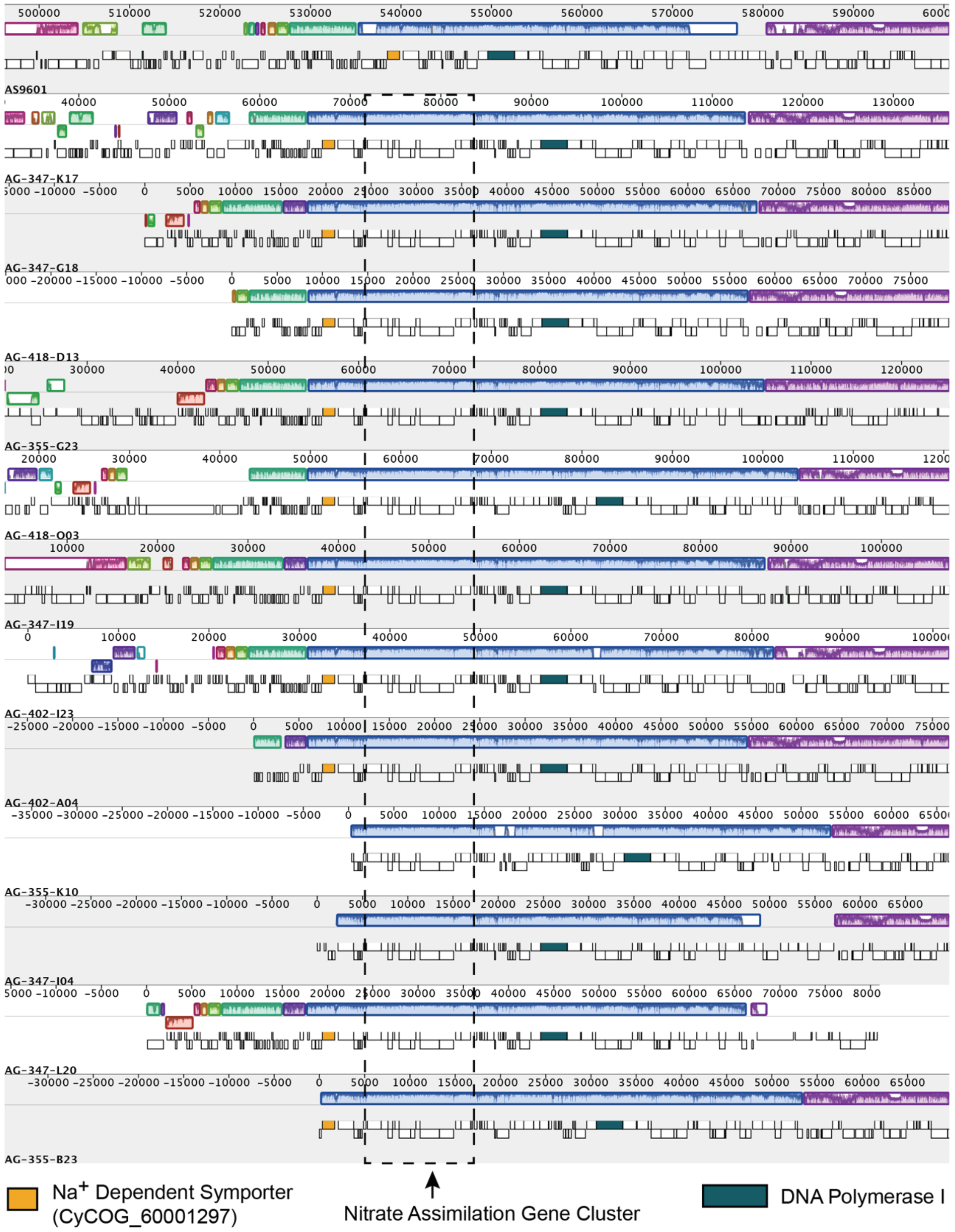


**Supplementary Fig. 3.** Mauve alignments of representative contigs from HLII *Prochlorococcus* single cell genome assemblies in comparison to the reference genome *Prochlorococcus* AS9601 (HLII; non-nitrate assimilating). The nitrate assimilation gene cluster is found in a local collinear block shared with AS9601. CyCOG_60001297 and DNA polymerase I genes are marked as reference points.

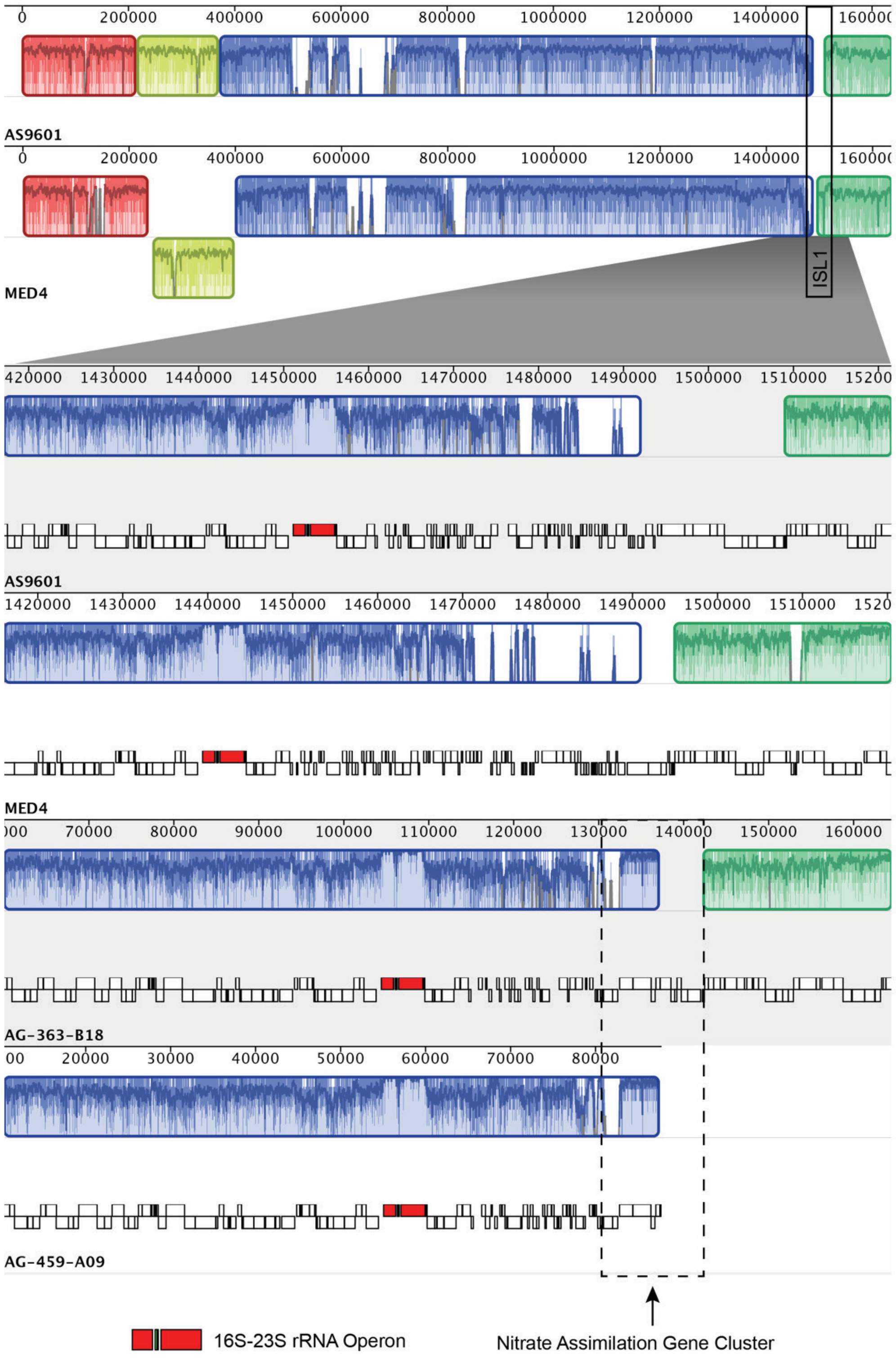


**Supplementary Fig. 4.** Mauve alignments of representative contigs from HLI *Prochlorococcus* single cell genome assemblies in comparison to the reference genomes *Prochlorococcus* MED4 (HLI) and *Prochlorococcus* AS9601 (HLII). The reference genomes do not contain the nitrate assimilation gene cluster. In single cells, this cluster is found in the genomic island ISL1 (sensu Kettler et al. 2007).

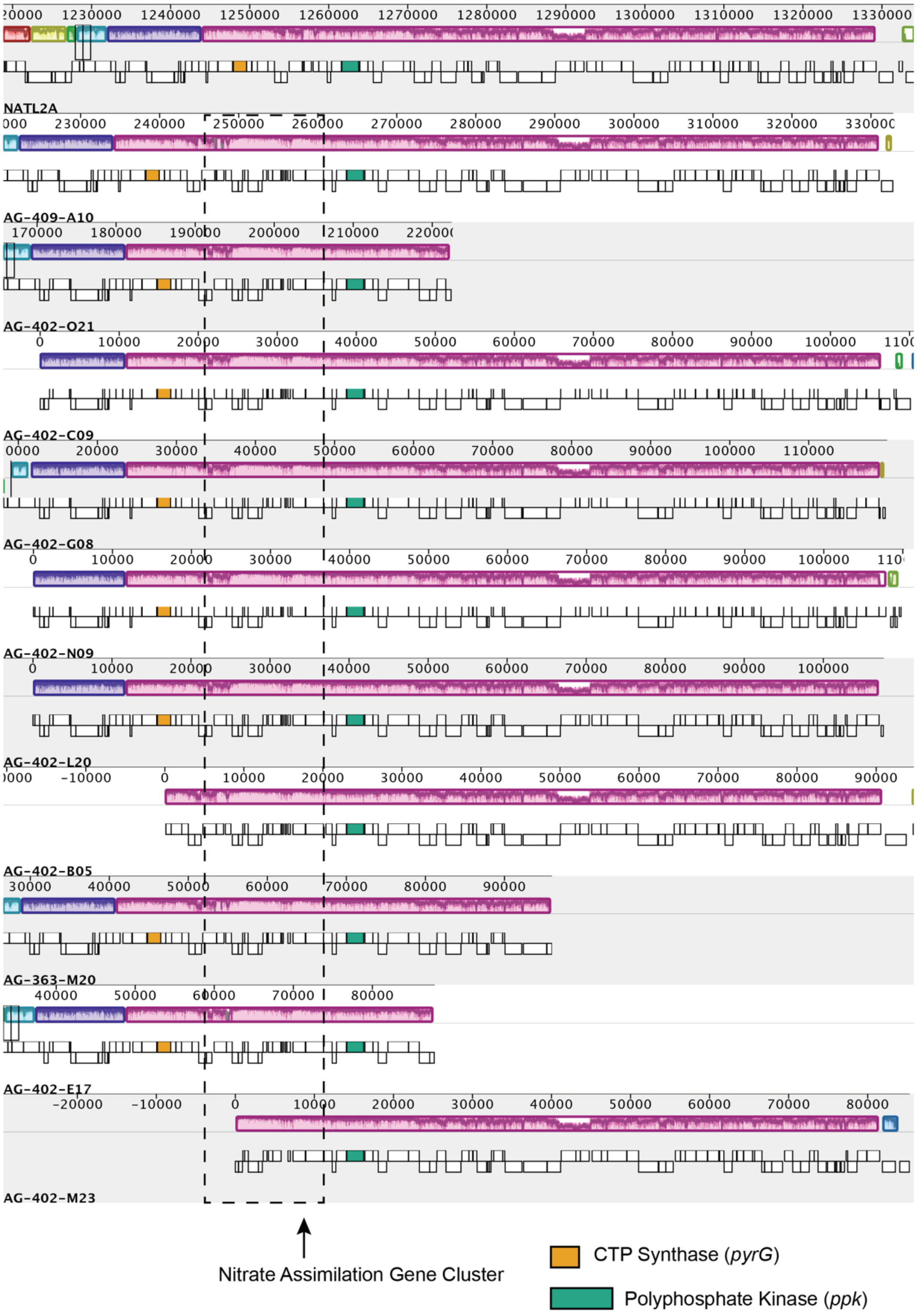


**Supplementary Fig. 5.** Mauve alignments of representative contigs from LLI *Prochlorococcus* single cell genome assemblies in comparison to the reference genome *Prochlorococcus* NATL2A (LLI; nitrite assimilation only). The nitrate assimilation gene cluster is found in a local collinear block shared with NATL2A. The *pyrG* and *ppk* genes are marked as reference points.

### Supplementary Information Tables

**Supplementary Table 1.** Estimates of dN/dS and results of tests for adaptive evolution among codon sites for *Prochlorococcus* and *Synechococcus*. Bolded LRT statistic values are chi-square critical values that meet a significance level of <0.001. For all genes, the inclusion of a class of neutral sites (M1) fits the data better than one dN/dS value for all sites (M0). While the inclusion of a class of sites under positive selection may be statistically justified under the M2 and M8 models, all dN/dS values are well below 1 suggesting that most sites are under purifying or neutral selection.

| | | log-likelihood of site models for adaptive evolution | | | | | likelihood ratio test (LRT) statistic for model pairs (degrees of freedom) | | |
|---|---|---|---|---|---|---|---|---|---|
| **gene** | **dN/dS** | **M0** | **M1** | **M2** | **M7** | **M8** | **M0 vs. M1 (1)** | **M1 vs. M2 (2)** | **M7 vs. M8 (2)** |
| ***gyrB*** | 0.036 | -105998 | -101367 | -101367 | -102434 | -100472 | **9263** | 0 | **3924** |
| ***pstB*** | 0.045 | -39945 | -39466 | -39466 | -38543 | -38484 | **957** | 0 | **118** |
| ***amtB*** | 0.066 | -70319 | -68351 | -68234 | -68226 | -67620 | **3935** | **235** | **1212** |
| ***glnA*** | 0.031 | -66948 | -65905 | -65905 | -65462 | -65118 | **2086** | 0 | **688** |
| ***glsF*** | 0.103 | -267696 | -249591 | -249591 | -253085 | -246724 | **36210** | 0 | **12722** |
| ***napA*** | 0.054 | -23709 | -23028 | -23028 | -23201 | -22961 | **1363** | 0 | **480** |
| ***narB*** | 0.119 | -40791 | -38790 | -38790 | -39102 | -38498 | **4001** | 0 | **1209** |
| ***moaA*** | 0.206 | -24794 | -22973 | -22931 | -23268 | -22770 | **3643** | **84** | **996** |
| ***focA*** | 0.078 | -18477 | -17701 | -17701 | -17895 | -17615 | **1553** | 0 | **562** |
| ***nirA*** | 0.109 | -54829 | -52361 | -52361 | -52063 | -51470 | **4937** | 0 | **1187** |

**Supplementary Table 2.** Tests for adaptive evolution among codon sites for the foreground HLII branch. Background lineages include *Synechococcus* and all other *Prochlorococcus*. The null model (H0) allows dN/dS to vary among sites and includes a fixed class of neutral sites in both the foreground and background branches. H1 allows classes of sites with dN/dS > 1 in the foreground branch only. Bolded LRT statistic values are chi-square critical values that meet a significance level of <0.001. While the inclusion of sites under positive selection is justified for the *gyrB* and *nirA* genes, most sites appear to be under purifying selection.

| gene | hypothesis log-likelihood | | site class 0 | site class 1 | site class 2a | site class 2b | LRT statistic |
|---|---|---|---|---|---|---|---|
| ***gyrB*** | H0 -104105 | proportion of sites | 0.98759 | 0.00138 | 0.01102 | 0.00002 | **6074** |
| | | background dN/dS | 0 | 1 | 0 | 1 | |
| | | foreground dN/dS | 0 | 1 | 1 | 1 | |
| | H1 -101068 | proportion of sites | 0.99666 | 0.00177 | 0.00157 | 0 | |
| | | background dN/dS | 0.00908 | 1 | 0.00908 | 1 | |
| | | foreground dN/dS | 0.00908 | 1 | 1 | 1 | |
| ***pstB*** | H0 -39463 | proportion of sites | 0.99801 | 0.00161 | 0.00038 | 0 | 0 |
| | | background dN/dS | 0.03255 | 1 | 0.03255 | 1 | |
| | | foreground dN/dS | 0.03255 | 1 | 1 | 1 | |
| | H1 -39463 | proportion of sites | 0.99801 | 0.00161 | 0.00038 | 0 | |
| | | background dN/dS | 0.03255 | 1 | 0.03255 | 1 | |
| | | foreground dN/dS | 0.03255 | 1 | 1 | 1 | |
| ***amtB*** | H0 -68231 | proportion of sites | 0.9926 | 0.00663 | 0.00076 | 0.00001 | -2 |
| | | background dN/dS | 0.02052 | 1 | 0.02052 | 1 | |
| | | foreground dN/dS | 0.02052 | 1 | 1 | 1 | |
| | H1 -68232 | proportion of sites | 0.99288 | 0.00657 | 0.00054 | 0 | |
| | | background dN/dS | 0.0206 | 1 | 0.0206 | 1 | |
| | | foreground dN/dS | 0.0206 | 1 | 2.17216 | 2.17216 | |
| ***glnA*** | H0 -65896 | proportion of sites | 0.99839 | 0.00144 | 0.00018 | 0 | 0 |
| | | background dN/dS | 0.01722 | 1 | 0.01722 | 1 | |
| | | foreground dN/dS | 0.01722 | 1 | 1 | 1 | |
| | H1 -65896 | proportion of sites | 0.99839 | 0.00144 | 0.00018 | 0 | |
| | | background dN/dS | 0.01722 | 1 | 0.01722 | 1 | |
| | | foreground dN/dS | 0.01722 | 1 | 1 | 1 | |
| ***glsF*** | H0 -249282 | proportion of sites | 0.99092 | 0.00679 | 0.00227 | 0.00002 | 0 |
| | | background dN/dS | 0.0227 | 1 | 0.0227 | 1 | |
| | | foreground dN/dS | 0.0227 | 1 | 1 | 1 | |
| | H1 -249282 | proportion of sites | 0.99092 | 0.00679 | 0.00227 | 0.00002 | |
| | | background dN/dS | 0.0227 | 1 | 0.0227 | 1 | |
| | | foreground dN/dS | 0.0227 | 1 | 1 | 1 | |
| ***napA*** | H0 -23009 | proportion of sites | 0.97966 | 0.01219 | 0.00805 | 0.0001 | 0 |
| | | background dN/dS | 0.00928 | 1 | 0.00928 | 1 | |
| | | foreground dN/dS | 0.00928 | 1 | 1 | 1 | |
| | H1 -23009 | proportion of sites | 0.97966 | 0.01219 | 0.00805 | 0.0001 | |
| | | background dN/dS | 0.00928 | 1 | 0.00928 | 1 | |
| | | foreground dN/dS | 0.00928 | 1 | 1 | 1 | |
| ***narB*** | H0 -38711 | proportion of sites | 0.94311 | 0.02164 | 0.03446 | 0.00079 | 0 |
| | | background dN/dS | 0.02489 | 1 | 0.02489 | 1 | |
| | | foreground dN/dS | 0.02489 | 1 | 1 | 1 | |
| | H1 -38711 | proportion of sites | 0.94311 | 0.02164 | 0.03447 | 0.00079 | |
| | | background dN/dS | 0.02489 | 1 | 0.02489 | 1 | |
| | | foreground dN/dS | 0.02489 | 1 | 1 | 1 | |
| ***moaA*** | H0 -22958 | proportion of sites | 0.94068 | 0.03375 | 0.02468 | 0.00089 | 0 |
| | | background dN/dS | 0.03882 | 1 | 0.03882 | 1 | |
| | | foreground dN/dS | 0.03882 | 1 | 1 | 1 | |
| | H1 -22958 | proportion of sites | 0.94068 | 0.03375 | 0.02468 | 0.00089 | |
| | | background dN/dS | 0.03882 | 1 | 0.03882 | 1 | |
| | | foreground dN/dS | 0.03882 | 1 | 1 | 1 | |
| ***nirA*** | H0 -52348 | proportion | 0.97996 | 0.01302 | 0.00693 | 0.00009 | **14** |
| | | background dN/dS | 0.04327 | 1 | 0.04327 | 1 | |
| | | foreground dN/dS | 0.04327 | 1 | 1 | 1 | |
| | H1 -52341 | proportion | 0.97254 | 0.0124 | 0.01487 | 0.00019 | |
| | | background dN/dS | 0.04293 | 1 | 0.04293 | 1 | |
| | | foreground dN/dS | 0.04293 | 1 | 1 | 1 | |

**Supplementary Table 3.** Tests for adaptive evolution among codon sites for the foreground LLI branch. Background lineages include *Synechococcus* and all other *Prochlorococcus*. While the inclusion of sites under positive selection is justified for the *gyrB* and *narB* genes, most sites appear to be under purifying selection. See Supplementary Table 2 for model descriptions.

| gene | hypothesis log-likelihood | | site class 0 | site class 1 | site class 2a | site class 2b | LRT statistic |
|---|---|---|---|---|---|---|---|
| ***gyrB*** | H0 -101367 | proportion of sites | 0.99813 | 0.00187 | 0 | 0 | **22** |
| | | background dN/dS | 0.01062 | 1 | 0.01062 | 1 | |
| | | foreground dN/dS | 0.01062 | 1 | 1 | 1 | |
| | H1 -101356 | proportion of sites | 0.99768 | 0.00186 | 0.00046 | 0 | |
| | | background dN/dS | 0.01053 | 1 | 0.01053 | 1 | |
| | | foreground dN/dS | 0.01053 | 1 | 1 | 1 | |
| ***pstB*** | H0 -39407 | proportion of sites | 0.98046 | 0.00147 | 0.01805 | 0.00003 | 0 |
| | | background dN/dS | 0.03172 | 1 | 0.03172 | 1 | |
| | | foreground dN/dS | 0.03172 | 1 | 1 | 1 | |
| | H1 -39407 | proportion of sites | 0.98046 | 0.00147 | 0.01805 | 0.00003 | |
| | | background dN/dS | 0.03172 | 1 | 0.03172 | 1 | |
| | | foreground dN/dS | 0.03172 | 1 | 1 | 1 | |
| ***amtB*** | H0 -68211 | proportion of sites | 0.99157 | 0.00673 | 0.00169 | 0.00001 | 0 |
| | | background dN/dS | 0.01992 | 1 | 0.01992 | 1 | |
| | | foreground dN/dS | 0.01992 | 1 | 1 | 1 | |
| | H1 -68211 | proportion of sites | 0.9918 | 0.00677 | 0.00143 | 0.00001 | |
| | | background dN/dS | 0.01992 | 1 | 0.01992 | 1 | |
| | | foreground dN/dS | 0.01992 | 1 | 1 | 1 | |
| ***glnA*** | H0 -65885 | proportion of sites | 0.99799 | 0.00141 | 0.0006 | 0 | 0 |
| | | background dN/dS | 0.01727 | 1 | 0.01727 | 1 | |
| | | foreground dN/dS | 0.01727 | 1 | 1 | 1 | |
| | H1 -65885 | proportion of sites | 0.99799 | 0.00141 | 0.0006 | 0 | |
| | | background dN/dS | 0.01727 | 1 | 0.01727 | 1 | |
| | | foreground dN/dS | 0.01727 | 1 | 1 | 1 | |
| ***glsF*** | H0 -249371 | proportion of sites | 0.98865 | 0.00686 | 0.00445 | 0.00003 | 0 |
| | | background dN/dS | 0.02306 | 1 | 0.02306 | 1 | |
| | | foreground dN/dS | 0.02306 | 1 | 1 | 1 | |
| | H1 -249371 | proportion of sites | 0.98865 | 0.00686 | 0.00445 | 0.00003 | |
| | | background dN/dS | 0.02306 | 1 | 0.02306 | 1 | |
| | | foreground dN/dS | 0.02306 | 1 | 1 | 1 | |
| ***napA*** | H0 -23014 | proportion of sites | 0.97128 | 0.01248 | 0.01604 | 0.00021 | 0 |
| | | background dN/dS | 0.01004 | 1 | 0.01004 | 1 | |
| | | foreground dN/dS | 0.01004 | 1 | 1 | 1 | |
| | H1 -23014 | proportion of sites | 0.97122 | 0.01246 | 0.01611 | 0.00021 | |
| | | background dN/dS | 0.01004 | 1 | 0.01004 | 1 | |
| | | foreground dN/dS | 0.01004 | 1 | 1 | 1 | |
| ***narB*** | H0 -38790 | proportion of sites | 0.97596 | 0.02404 | 0 | 0 | **40** |
| | | background dN/dS | 0.02939 | 1 | 0.02939 | 1 | |
| | | foreground dN/dS | 0.02939 | 1 | 1 | 1 | |
| | H1 -38770 | proportion of sites | 0.9577 | 0.02346 | 0.01839 | 0.00045 | |
| | | background dN/dS | 0.02762 | 1 | 0.02762 | 1 | |
| | | foreground dN/dS | 0.02762 | 1 | 1 | 1 | |
| ***moaA*** | H0 -22961 | proportion of sites | 0.90692 | 0.03565 | 0.05526 | 0.00217 | 0 |
| | | background dN/dS | 0.03704 | 1 | 0.03704 | 1 | |
| | | foreground dN/dS | 0.03704 | 1 | 1 | 1 | |
| | H1 -22961 | proportion of sites | 0.90692 | 0.03565 | 0.05526 | 0.00217 | |
| | | background dN/dS | 0.03704 | 1 | 0.03704 | 1 | |
| | | foreground dN/dS | 0.03704 | 1 | 1 | 1 | |
| ***focA*** | H0 -17695 | proportion of sites | 0.97605 | 0.01411 | 0.00969 | 0.00014 | 0 |
| | | background dN/dS | 0.01425 | 1 | 0.01425 | 1 | |
| | | foreground dN/dS | 0.01425 | 1 | 1 | 1 | |
| | H1 -17695 | proportion of sites | 0.97605 | 0.01411 | 0.0097 | 0.00014 | |
| | | background dN/dS | 0.01425 | 1 | 0.01425 | 1 | |
| | | foreground dN/dS | 0.01425 | 1 | 1 | 1 | |
| ***nirA* type I** | H0 -52349 | proportion of sites | 0.97292 | 0.01243 | 0.01446 | 0.00018 | 0 |
| | | background dN/dS | 0.04333 | 1 | 0.04333 | 1 | |
| | | foreground dN/dS | 0.04333 | 1 | 1 | 1 | |
| | H1 -52349 | proportion of sites | 0.97292 | 0.01243 | 0.01446 | 0.00018 | |
| | | background dN/dS | 0.04333 | 1 | 0.04333 | 1 | |
| | | foreground dN/dS | 0.04333 | 1 | 1 | 1 | |
| ***nirA* type II** | H0 -52358 | proportion of sites | 0.97139 | 0.01216 | 0.01625 | 0.0002 | 0 |
| | | background dN/dS | 0.0442 | 1 | 0.0442 | 1 | |
| | | foreground dN/dS | 0.0442 | 1 | 1 | 1 | |
| | H1 -52358 | proportion of sites | 0.97139 | 0.01216 | 0.01625 | 0.0002 | |
| | | background dN/dS | 0.0442 | 1 | 0.0442 | 1 | |
| | | foreground dN/dS | 0.0442 | 1 | 1 | 1 | |

**Supplementary Table 4.** Genomes and associated IMG accession numbers in the final data set. Clade indicates the respective *Synechococcus* subcluster or *Prochlorococcus* clade. Results from the *narB* PCR screening assay are presented as a binary (0 = negative; 1 = positive; n.d = not determined). Estimated genome recovery was determined using checkM (Parks et al. 2015) and the presence/absence of annotated reductase and transporter genes for nitrite (*nirA* and *focA*) and nitrate (*narB* and *napA*) in the assembly are as given as a binary (0 = absent; 1 = present).

| Genome | IMG ID | Clade | *narB* PCR | % Genome Recovery | *nirA* | *focA* | *narB* | *napA* |
|---|---|---|---|---|---|---|---|---|
| RCC307 | 2623620283 | Syn 5.3 | n.d. | >99 | 1 | 1 | 1 | 1 |
| BL107 | 2623620351 | Syn 5.1 | n.d. | >99 | 1 | 1 | 1 | 1 |
| CC9311 | 2623620876 | Syn 5.1 | n.d. | >99 | 1 | 1 | 1 | 1 |
| CC9605 | 2606217400 | Syn 5.1 | n.d. | >99 | 1 | 1 | 1 | 1 |
| CC9616 | 2517093019 | Syn 5.1 | n.d. | >99 | 1 | 0 | 1 | 1 |
| CC9902 | 2606217272 | Syn 5.1 | n.d. | >99 | 1 | 1 | 1 | 1 |
| KORDI-100 | 2507262013 | Syn 5.1 | n.d. | >99 | 1 | 0 | 1 | 1 |
| KORDI-49 | 2507262011 | Syn 5.1 | n.d. | >99 | 1 | 1 | 1 | 1 |
| KORDI-52 | 2507262012 | Syn 5.1 | n.d. | >99 | 1 | 1 | 1 | 1 |
| MIT9504 | 2681812951 | Syn 5.1 | n.d. | >99 | 1 | 1 | 1 | 1 |
| MIT9508 | 2681812952 | Syn 5.1 | n.d. | >99 | 1 | 1 | 1 | 1 |
| MIT9509 | 2681812953 | Syn 5.1 | n.d. | >99 | 1 | 1 | 1 | 1 |
| RS9916 | 2623620281 | Syn 5.1 | n.d. | >99 | 1 | 1 | 1 | 1 |
| RS9917 | 638341213 | Syn 5.1 | n.d. | >99 | 1 | 1 | 0 | 0 |
| WH7803 | 2623620330 | Syn 5.1 | n.d. | >99 | 1 | 1 | 1 | 1 |
| WH7805 | 2623620868 | Syn 5.1 | n.d. | >99 | 1 | 1 | 1 | 1 |
| WH8016 | 2507262052 | Syn 5.1 | n.d. | >99 | 1 | 1 | 1 | 1 |
| WH8020 | 2645727663 | Syn 5.1 | n.d. | >99 | 1 | 1 | 1 | 1 |
| WH8102 | 2606217514 | Syn 5.1 | n.d. | >99 | 1 | 0 | 1 | 1 |
| WH8103 | 2687453380 | Syn 5.1 | n.d. | >99 | 1 | 0 | 1 | 1 |
| WH8109 | 2563366603 | Syn 5.1 | n.d. | >99 | 1 | 1 | 1 | 1 |
| AG-679-C18 | 2716884671 | Syn 5.1 | n.d. | 87.57 | 1 | 1 | 1 | 1 |
| AG-670-B23 | 2716884610 | Syn 5.1 | n.d. | 86.46 | 1 | 1 | 1 | 1 |
| AG-673-D02 | 2716884257 | Syn 5.1 | n.d. | 86.21 | 1 | 1 | 1 | 1 |
| AG-670-A05 | 2716884608 | Syn 5.1 | n.d. | 81.39 | 1 | 0 | 1 | 1 |
| AG-679-B05 | 2716884670 | Syn 5.1 | n.d. | 81.1 | 0 | 0 | 1 | 1 |
| AG-673-F03 | 2716884618 | Syn 5.1 | n.d. | 77.9 | 1 | 1 | 1 | 1 |
| AG-670-F22 | 2716884613 | Syn 5.1 | n.d. | 73.1 | 1 | 1 | 1 | 1 |
| AG-679-D13 | 2716884624 | Syn 5.1 | n.d. | 72.55 | 1 | 1 | 1 | 1 |
| AG-673-A03 | 2716884615 | Syn 5.1 | n.d. | 69.77 | 0 | 0 | 1 | 1 |
| AG-676-E23 | 2716884621 | Syn 5.1 | n.d. | 67.53 | 1 | 1 | 1 | 1 |
| AG-676-C06 | 2716884620 | Syn 5.1 | n.d. | 66.44 | 1 | 1 | 1 | 1 |
| AG-679-A02 | 2716884668 | Syn 5.1 | n.d. | 62.76 | 1 | 1 | 1 | 1 |
| AG-683-B21 | 2716884674 | Syn 5.1 | n.d. | 61.67 | 1 | 1 | 1 | 1 |
| AG-673-A10 | 2716884616 | Syn 5.1 | n.d. | 58.11 | 1 | 1 | 1 | 0 |
| AG-673-B08 | 2716884617 | Syn 5.1 | n.d. | 52.05 | 1 | 1 | 1 | 1 |
| AG-683-A03 | 2716884626 | Syn 5.1 | n.d. | 52.04 | 1 | 1 | 1 | 1 |
| AG-670-D07 | 2716884611 | Syn 5.1 | n.d. | 51.98 | 1 | 0 | 1 | 1 |
| AG-683-G11 | 2716884675 | Syn 5.1 | n.d. | 50.17 | 1 | 1 | 1 | 1 |
| AG-670-A19 | 2716884609 | Syn 5.1 | n.d. | 47.37 | 1 | 1 | 1 | 1 |
| AG-670-A04 | 2716884607 | Syn 5.1 | n.d. | 45.29 | 1 | 1 | 1 | 1 |
| AG-402-L18 | 2667527345 | HL.OTHER | 0 | 87.09 | 0 | 0 | 0 | 0 |
| EQPAC1 | 2606217689 | HLI | n.d. | >99 | 0 | 0 | 0 | 0 |
| MED4 | 2606217259 | HLI | n.d. | >99 | 0 | 0 | 0 | 0 |
| MIT9515 | 2623620345 | HLI | n.d. | >99 | 0 | 0 | 0 | 0 |
| AG-418-M08 | 2716884409 | HLI | 0 | 98.51 | 0 | 0 | 0 | 0 |
| AG-402-N10 | 2667527275 | HLI | 0 | 95.11 | 0 | 0 | 0 | 0 |
| AG-347-J22 | 2667527322 | HLI | 0 | 89.27 | 0 | 0 | 0 | 0 |
| AG-459-A09 | 2716884789 | HLI | 1 | 84.74 | 1 | 0 | 1 | 1 |

| | | | | | | | | |
|---|---|---|---|---|---|---|---|---|
| AG-459-O03 | 2716884798 | HLI | 0 | 72.14 | 0 | 0 | 0 | 0 |
| AG-459-P02 | 2716884459 | HLI | 1 | 59.92 | 1 | 0 | 1 | 1 |
| AG-459-M13 | 2716884796 | HLI | 1 | 59.24 | 0 | 0 | 0 | 0 |
| AG-459-P16 | 2716884460 | HLI | 0 | 57.02 | 0 | 0 | 0 | 0 |
| AG-459-J20 | 2716884795 | HLI | 1 | 50.32 | 1 | 0 | 1 | 1 |
| AG-335-D02 | 2716884715 | HLI | n.d. | 45.83 | 1 | 0 | 1 | 1 |
| AG-335-O19 | 2716884722 | HLI | n.d. | 44.14 | 1 | 0 | 1 | 1 |
| AG-459-A01 | 2716884456 | HLI | 0 | 41.54 | 0 | 0 | 0 | 0 |
| AG-459-P19 | 2716884800 | HLI | 0 | 39.27 | 0 | 0 | 0 | 0 |
| AS9601 | 2623620959 | HLII | n.d. | >99 | 0 | 0 | 0 | 0 |
| GP2 | 2606217606 | HLII | n.d. | >99 | 0 | 0 | 0 | 0 |
| MIT0604 | 2606217688 | HLII | n.d. | >99 | 1 | 0 | 1 | 1 |
| MIT1314 | 2681813573 | HLII | n.d. | >99 | 0 | 0 | 0 | 0 |
| MIT9107 | 2606217692 | HLII | n.d. | >99 | 0 | 0 | 0 | 0 |
| MIT9116 | 2606217690 | HLII | n.d. | >99 | 0 | 0 | 0 | 0 |
| MIT9123 | 2606217318 | HLII | n.d. | >99 | 0 | 0 | 0 | 0 |
| MIT9201 | 2606217687 | HLII | n.d. | >99 | 0 | 0 | 0 | 0 |
| MIT9202 | 2623620984 | HLII | n.d. | >99 | 0 | 0 | 0 | 0 |
| MIT9215 | 2606217559 | HLII | n.d. | >99 | 0 | 0 | 0 | 0 |
| MIT9301 | 2623620961 | HLII | 0 | >99 | 0 | 0 | 0 | 0 |
| MIT9302 | 2606217691 | HLII | n.d. | >99 | 0 | 0 | 0 | 0 |
| MIT9311 | 2606217680 | HLII | n.d. | >99 | 0 | 0 | 0 | 0 |
| MIT9312 | 2606217708 | HLII | n.d. | >99 | 0 | 0 | 0 | 0 |
| MIT9314 | 2606217312 | HLII | n.d. | >99 | 0 | 0 | 0 | 0 |
| MIT9321 | 2606217683 | HLII | n.d. | >99 | 0 | 0 | 0 | 0 |
| MIT9322 | 2606217679 | HLII | n.d. | >99 | 0 | 0 | 0 | 0 |
| MIT9401 | 2606217316 | HLII | n.d. | >99 | 0 | 0 | 0 | 0 |
| SB | 2606217677 | HLII | 1 | >99 | 1 | 0 | 1 | 1 |
| AG-402-K21 | 2667527344 | HLII | 0 | 98.1 | 0 | 0 | 0 | 0 |
| AG-402-M15 | 2667527272 | HLII | 0 | 97.28 | 0 | 0 | 0 | 0 |
| AG-412-J13 | 2716884760 | HLII | n.d. | 97.11 | 1 | 0 | 1 | 1 |
| AG-355-N16 | 2667527304 | HLII | 0 | 96.33 | 0 | 0 | 0 | 0 |
| AG-402-L23 | 2667527271 | HLII | 1 | 95.79 | 1 | 0 | 1 | 1 |
| AG-355-B18 | 2667527285 | HLII | 0 | 95.7 | 0 | 0 | 0 | 0 |
| AG-347-L20 | 2667527248 | HLII | 1 | 95.24 | 1 | 0 | 1 | 1 |
| AG-402-N08 | 2716884383 | HLII | 1 | 95.15 | 1 | 0 | 1 | 1 |
| AG-347-G18 | 2667527329 | HLII | 1 | 95.03 | 1 | 0 | 1 | 1 |
| AG-355-M02 | 2667527301 | HLII | 1 | 94.88 | 0 | 0 | 0 | 0 |
| AG-355-O17 | 2667527308 | HLII | 0 | 94.61 | 1 | 0 | 0 | 0 |
| AG-347-K16 | 2667527326 | HLII | 0 | 94.47 | 0 | 0 | 0 | 0 |
| AG-347-L02 | 2667527245 | HLII | 0 | 94.2 | 0 | 0 | 0 | 0 |
| AG-355-I04 | 2667527288 | HLII | 0 | 94.02 | 0 | 0 | 0 | 0 |
| AG-347-K18 | 2667527240 | HLII | 0 | 93.93 | 0 | 0 | 0 | 0 |
| AG-418-D13 | 2716884768 | HLII | 1 | 93.93 | 1 | 0 | 1 | 1 |
| AG-347-L17 | 2667527246 | HLII | 1 | 93.8 | 1 | 0 | 0 | 0 |
| AG-347-I06 | 2667527332 | HLII | 0 | 93.75 | 0 | 0 | 0 | 0 |
| AG-347-J19 | 2667527339 | HLII | 0 | 93.75 | 0 | 0 | 0 | 0 |
| AG-355-K20 | 2667527295 | HLII | 0 | 93.7 | 0 | 0 | 0 | 0 |
| AG-347-I19 | 2667527334 | HLII | 1 | 93.34 | 1 | 0 | 1 | 1 |
| AG-347-I21 | 2667527335 | HLII | 1 | 93.21 | 1 | 0 | 1 | 1 |
| AG-355-P11 | 2667527349 | HLII | 0 | 93.1 | 0 | 0 | 0 | 0 |
| AG-347-J21 | 2667527341 | HLII | 0 | 92.84 | 0 | 0 | 0 | 0 |
| AG-402-O16 | 2667527278 | HLII | 0 | 92.8 | 0 | 0 | 0 | 0 |
| AG-418-I20 | 2716884403 | HLII | 0 | 92.66 | 0 | 0 | 0 | 0 |
| AG-355-P07 | 2667527348 | HLII | 0 | 92.53 | 0 | 0 | 0 | 0 |
| AG-402-I23 | 2667527263 | HLII | 1 | 92.53 | 1 | 0 | 1 | 1 |
| AG-442-N07 | 2716884446 | HLII | n.d. | 92.12 | 1 | 0 | 1 | 1 |
| AG-347-K19 | 2667527241 | HLII | 0 | 92.07 | 0 | 0 | 0 | 0 |
| AG-355-A09 | 2667527284 | HLII | 0 | 92.03 | 0 | 0 | 0 | 0 |

| | | | | | | | | |
|---|---|---|---|---|---|---|---|---|
| AG-355-J09 | 2667527291 | HLII | 1 | 91.91 | 1 | 0 | 1 | 1 |
| AG-355-P16 | 2667527351 | HLII | 0 | 91.85 | 1 | 0 | 0 | 0 |
| AG-347-K17 | 2667527239 | HLII | 1 | 91.53 | 1 | 0 | 1 | 1 |
| AG-347-M08 | 2716884361 | HLII | 1 | 91.53 | 0 | 0 | 0 | 0 |
| AG-347-J14 | 2667527338 | HLII | 0 | 91.17 | 0 | 0 | 0 | 0 |
| AG-355-N02 | 2667527303 | HLII | 0 | 91.17 | 0 | 0 | 0 | 0 |
| AG-355-N22 | 2667527306 | HLII | 0 | 90.85 | 0 | 0 | 0 | 0 |
| AG-347-J20 | 2667527340 | HLII | 0 | 90.67 | 0 | 0 | 0 | 0 |
| AG-347-K20 | 2667527242 | HLII | 0 | 90.49 | 0 | 0 | 0 | 0 |
| AG-347-E23 | 2716884358 | HLII | 1 | 90.22 | 0 | 0 | 0 | 0 |
| AG-347-I04 | 2716884359 | HLII | 1 | 90.04 | 1 | 0 | 1 | 1 |
| AG-347-M23 | 2667527252 | HLII | 1 | 89.58 | 1 | 0 | 1 | 1 |
| AG-347-G20 | 2667527330 | HLII | 0 | 89.4 | 1 | 0 | 1 | 1 |
| AG-418-P13 | 2716884414 | HLII | 0 | 89.4 | 1 | 0 | 1 | 1 |
| AG-402-K16 | 2667527343 | HLII | 1 | 89.18 | 1 | 0 | 1 | 1 |
| AG-347-K02 | 2667527324 | HLII | 1 | 89.09 | 0 | 0 | 0 | 0 |
| AG-355-N18 | 2667527347 | HLII | 1 | 88.72 | 1 | 0 | 1 | 1 |
| AG-355-P18 | 2667527352 | HLII | 0 | 88.68 | 0 | 0 | 0 | 0 |
| AG-402-F05 | 2716884380 | HLII | 1 | 88.67 | 1 | 0 | 1 | 1 |
| AG-347-N19 | 2667527253 | HLII | 1 | 87.77 | 0 | 0 | 0 | 0 |
| AG-355-J23 | 2667527292 | HLII | 0 | 87.41 | 0 | 0 | 0 | 0 |
| AG-347-I15 | 2667527333 | HLII | 0 | 86.96 | 0 | 0 | 0 | 0 |
| AG-347-O22 | 2667527255 | HLII | 1 | 86.91 | 1 | 0 | 1 | 1 |
| AG-355-L02 | 2667527297 | HLII | 0 | 86.87 | 0 | 0 | 0 | 0 |
| AG-347-J06 | 2716884360 | HLII | 1 | 86.85 | 1 | 0 | 1 | 1 |
| AG-402-K22 | 2667527267 | HLII | 1 | 86.19 | 0 | 0 | 1 | 1 |
| AG-347-E03 | 2716884357 | HLII | 1 | 86.01 | 0 | 0 | 0 | 0 |
| AG-347-M18 | 2667527251 | HLII | 0 | 85.78 | 0 | 0 | 0 | 0 |
| AG-347-K10 | 2716884262 | HLII | 1 | 85.64 | 0 | 0 | 0 | 0 |
| AG-355-K15 | 2667527294 | HLII | 0 | 85.39 | 0 | 0 | 0 | 0 |
| AG-355-L20 | 2667527298 | HLII | 0 | 85.24 | 0 | 0 | 0 | 0 |
| AG-418-F08 | 2716884769 | HLII | 1 | 84.89 | 0 | 0 | 0 | 0 |
| AG-355-G23 | 2667527287 | HLII | 1 | 84.83 | 1 | 0 | 1 | 1 |
| AG-347-M15 | 2667527250 | HLII | 0 | 84.38 | 0 | 0 | 0 | 0 |
| AG-402-C22 | 2716884378 | HLII | 1 | 84.01 | 0 | 0 | 0 | 0 |
| AG-418-J17 | 2716884404 | HLII | 0 | 83.29 | 0 | 0 | 0 | 0 |
| AG-418-P06 | 2716884413 | HLII | 0 | 83.02 | 0 | 0 | 0 | 0 |
| AG-347-L19 | 2667527247 | HLII | 1 | 82.52 | 0 | 0 | 0 | 0 |
| AG-347-L21 | 2667527249 | HLII | 0 | 81.95 | 0 | 0 | 0 | 0 |
| AG-402-N23 | 2667527277 | HLII | 0 | 81.79 | 0 | 0 | 0 | 0 |
| AG-355-I20 | 2667527289 | HLII | 1 | 81.16 | 1 | 0 | 1 | 1 |
| AG-355-M18 | 2667527302 | HLII | 0 | 80.66 | 0 | 0 | 0 | 0 |
| AG-402-A04 | 2716884372 | HLII | 1 | 80.21 | 1 | 0 | 1 | 1 |
| AG-355-K03 | 2716884364 | HLII | 1 | 80.16 | 0 | 0 | 0 | 0 |
| AG-347-G22 | 2667527331 | HLII | 0 | 79.98 | 0 | 0 | 0 | 0 |
| AG-418-O03 | 2716884412 | HLII | 1 | 79.48 | 1 | 0 | 1 | 1 |
| AG-355-B23 | 2667527286 | HLII | 1 | 79.38 | 1 | 0 | 1 | 1 |
| AG-347-J23 | 2667527323 | HLII | 0 | 79.08 | 0 | 0 | 0 | 0 |
| AG-418-O02 | 2716884411 | HLII | 1 | 77.94 | 1 | 0 | 1 | 1 |
| AG-459-N19 | 2716884797 | HLII | 0 | 76.81 | 0 | 0 | 0 | 0 |
| AG-347-K15 | 2667527325 | HLII | 0 | 76.13 | 0 | 0 | 0 | 0 |
| AG-347-C10 | 2716884356 | HLII | 1 | 75.07 | 0 | 0 | 0 | 0 |
| AG-347-B23 | 2667527321 | HLII | 0 | 74.67 | 0 | 0 | 0 | 0 |
| AG-355-P15 | 2667527350 | HLII | 0 | 74.67 | 0 | 0 | 0 | 0 |
| AG-355-L22 | 2667527300 | HLII | 0 | 74.34 | 0 | 0 | 0 | 0 |
| AG-459-O09 | 2716884458 | HLII | 1 | 74.05 | 1 | 0 | 1 | 1 |
| AG-449-C14 | 2716884448 | HLII | n.d. | 72.82 | 1 | 0 | 1 | 1 |
| AG-418-G18 | 2716884770 | HLII | 1 | 72.54 | 0 | 0 | 0 | 0 |
| AG-355-J04 | 2667527290 | HLII | 0 | 71.95 | 0 | 0 | 0 | 0 |

| | | | | | | | | |
|---|---|---|---|---|---|---|---|---|
| AG-355-K23 | 2667527296 | HLII | 0 | 71.78 | 0 | 0 | 0 | 0 |
| AG-402-G23 | 2667527258 | HLII | 1 | 71.56 | 1 | 0 | 1 | 1 |
| AG-459-D04 | 2716884792 | HLII | 1 | 70.79 | 1 | 0 | 1 | 1 |
| AG-355-A18 | 2716884362 | HLII | 1 | 70.43 | 1 | 0 | 1 | 1 |
| AG-418-G23 | 2716884402 | HLII | 0 | 70.2 | 0 | 0 | 0 | 0 |
| AG-355-K13 | 2667527293 | HLII | 0 | 70.01 | 0 | 0 | 0 | 0 |
| AG-418-J19 | 2716884405 | HLII | 1 | 69.7 | 0 | 0 | 0 | 0 |
| AG-459-B06 | 2716884790 | HLII | 1 | 69.57 | 1 | 0 | 1 | 1 |
| AG-418-F16 | 2716884401 | HLII | 1 | 68.34 | 0 | 0 | 0 | 0 |
| AG-355-N23 | 2667527307 | HLII | 0 | 67.37 | 0 | 0 | 0 | 0 |
| AG-424-M03 | 2716884775 | HLII | n.d. | 67.1 | 1 | 0 | 1 | 1 |
| AG-418-I21 | 2716884771 | HLII | 0 | 67.07 | 0 | 0 | 0 | 0 |
| AG-459-E08 | 2716884793 | HLII | 0 | 66.94 | 0 | 0 | 0 | 0 |
| AG-459-J14 | 2716884794 | HLII | 1 | 66.03 | 1 | 0 | 1 | 1 |
| AG-347-N23 | 2667527254 | HLII | 0 | 64.4 | 0 | 0 | 0 | 0 |
| AG-347-I22 | 2667527336 | HLII | 1 | 64.31 | 1 | 0 | 1 | 1 |
| AG-402-N17 | 2716884735 | HLII | 1 | 64.04 | 1 | 0 | 1 | 1 |
| AG-418-B17 | 2716884765 | HLII | 0 | 63.5 | 0 | 0 | 0 | 0 |
| AG-347-I23 | 2667527337 | HLII | 0 | 61.94 | 0 | 0 | 0 | 0 |
| AG-347-L13 | 2716884264 | HLII | 1 | 61.87 | 1 | 0 | 1 | 1 |
| AG-459-A02 | 2716884457 | HLII | 1 | 57.49 | 1 | 0 | 1 | 1 |
| AG-347-K22 | 2716884263 | HLII | 1 | 56.9 | 1 | 0 | 1 | 1 |
| AG-355-K10 | 2716884365 | HLII | 1 | 55.17 | 1 | 0 | 1 | 1 |
| AG-670-M18 | 2716884479 | HLII | n.d. | 53.68 | 0 | 0 | 1 | 1 |
| AG-347-K23 | 2667527244 | HLII | 0 | 53.12 | 0 | 0 | 0 | 0 |
| AG-355-A02 | 2667527283 | HLII | 1 | 50.86 | 0 | 0 | 1 | 1 |
| AG-402-O23 | 2667527280 | HLII | 0 | 49.74 | 0 | 0 | 0 | 0 |
| AG-418-L19 | 2716884407 | HLII | 0 | 49.59 | 0 | 0 | 0 | 0 |
| AG-363-A05 | 2667527356 | HLII | 0 | 49.23 | 0 | 0 | 0 | 0 |
| AG-347-K21 | 2667527243 | HLII | 0 | 48.69 | 0 | 0 | 0 | 0 |
| AG-347-B08 | 2716884355 | HLII | 1 | 47.28 | 0 | 0 | 0 | 0 |
| AG-459-P20 | 2716884801 | HLII | 0 | 47.01 | 0 | 0 | 0 | 0 |
| AG-355-J21 | 2667527346 | HLII | 0 | 45.29 | 0 | 0 | 0 | 0 |
| AG-347-J05 | 2716884726 | HLII | 1 | 43.29 | 0 | 0 | 0 | 0 |
| AG-355-J17 | 2716884363 | HLII | 1 | 43.1 | 1 | 0 | 1 | 1 |
| AG-355-N21 | 2667527305 | HLII | 0 | 41.38 | 0 | 0 | 0 | 0 |
| AG-355-L21 | 2667527299 | HLII | 0 | 40.49 | 0 | 0 | 0 | 0 |
| AG-449-K21 | 2716884788 | HLII | n.d. | 39.95 | 0 | 0 | 1 | 1 |
| AG-355-O19 | 2716884366 | HLII | 1 | 37.32 | 1 | 0 | 1 | 1 |
| AG-429-C19 | 2716884422 | HLII | n.d. | 35.87 | 1 | 0 | 1 | 1 |
| AG-355-P23 | 2667527353 | HLII | 0 | 34.64 | 0 | 0 | 0 | 0 |
| AG-455-E04 | 2716884452 | HLII | n.d. | 31.21 | 1 | 0 | 1 | 0 |
| AG-335-I15 | 2716884260 | HLII | n.d. | 26.69 | 1 | 0 | 1 | 1 |
| AG-418-K17 | 2716884406 | HLII | 0 | 76.77 | 0 | 0 | 0 | 0 |
| AG-363-P06 | 2667527317 | HLVI | 0 | 99.18 | 1 | 1 | 0 | 0 |
| AG-402-K10 | 2667527266 | HLVI | 0 | 93.61 | 0 | 0 | 0 | 0 |
| AG-363-B18 | 2667527363 | HLVI | 1 | 93.39 | 1 | 0 | 1 | 1 |
| MIT1223 | 2681813568 | LL.MIT1223 | n.d. | >99 | 1 | 1 | 0 | 0 |
| AG-402-N21 | 2667527276 | LL.MIT1223 | 0 | 79.8 | 0 | 0 | 0 | 0 |
| MIT1300 | 2681813570 | LL.OTHER | n.d. | >99 | 1 | 1 | 0 | 0 |
| MIT1307 | 2681813572 | LL.OTHER | n.d. | >99 | 1 | 1 | 0 | 0 |
| MIT1341 | 2681813574 | LL.OTHER | n.d. | >99 | 1 | 1 | 0 | 0 |
| AG-363-P19 | 2667527320 | LL.OTHER | 0 | 94.02 | 1 | 1 | 0 | 0 |
| AG-363-A16 | 2667527359 | LL.OTHER | 0 | 93.34 | 1 | 1 | 0 | 0 |
| AG-363-K07 | 2667527371 | LL.OTHER | 0 | 91.21 | 1 | 1 | 0 | 0 |
| AG-363-C20 | 2667527366 | LL.OTHER | 0 | 90.22 | 0 | 0 | 0 | 0 |
| AG-363-P08 | 2667527318 | LL.OTHER | 0 | 85.14 | 1 | 1 | 0 | 0 |
| AG-363-N20 | 2667527311 | LL.OTHER | 0 | 77.85 | 1 | 1 | 0 | 0 |
| AG-363-O06 | 2667527312 | LL.OTHER | 0 | 77.58 | 1 | 1 | 0 | 0 |

| | | | | | | | | |
|---|---|---|---|---|---|---|---|---|
| AG-363-O15 | 2667527313 | LL.OTHER | 0 | 77.12 | 1 | 1 | 0 | 0 |
| AG-363-B04 | 2667527360 | LL.OTHER | 0 | 75.68 | 0 | 0 | 0 | 0 |
| AG-363-P01 | 2667527316 | LL.OTHER | 0 | 73.19 | 1 | 1 | 0 | 0 |
| AG-363-M17 | 2667527375 | LL.OTHER | 0 | 72.28 | 1 | 0 | 0 | 0 |
| AG-363-J23 | 2667527370 | LL.OTHER | 0 | 67.53 | 0 | 0 | 0 | 0 |
| AG-363-P15 | 2667527319 | LL.OTHER | 0 | 60.4 | 0 | 0 | 0 | 0 |
| AG-363-O16 | 2667527314 | LL.OTHER | 0 | 58.69 | 1 | 1 | 0 | 0 |
| AG-363-I04 | 2667527309 | LL.OTHER | 0 | 50.82 | 1 | 1 | 0 | 0 |
| AG-363-L02 | 2667527372 | LL.OTHER | 0 | 50.4 | 0 | 0 | 0 | 0 |
| AG-363-A06 | 2667527357 | LL.OTHER | 0 | 47.87 | 1 | 1 | 0 | 0 |
| AG-363-A15 | 2667527358 | LL.OTHER | 0 | 44.55 | 0 | 0 | 0 | 0 |
| AG-402-K05 | 2667527265 | LL.OTHER | 0 | 41.38 | 0 | 0 | 0 | 0 |
| AG-363-L19 | 2667527374 | LL.OTHER | 0 | 37.93 | 0 | 0 | 0 | 0 |
| AG-363-O21 | 2667527315 | LL.OTHER | 0 | 36.21 | 0 | 0 | 0 | 0 |
| MIT0801 | 2606217560 | LLI | n.d. | >99 | 1 | 1 | 0 | 0 |
| MIT0912 | 2681812899 | LLI | n.d. | >99 | 1 | 1 | 1 | 1 |
| MIT0913 | 2681812900 | LLI | n.d. | >99 | 1 | 1 | 1 | 1 |
| MIT0915 | 2681812901 | LLI | n.d. | >99 | 1 | 1 | 1 | 1 |
| MIT0917 | 2681812859 | LLI | 1 | >99 | 1 | 0 | 1 | 1 |
| MIT1013 | 2681812904 | LLI | n.d. | >99 | 1 | 1 | 0 | 0 |
| MIT1214 | 2681813567 | LLI | n.d. | >99 | 1 | 1 | 0 | 0 |
| NATL1A | 2623620348 | LLI | n.d. | >99 | 1 | 1 | 0 | 0 |
| NATL2A | 2606217240 | LLI | 0 | >99 | 1 | 1 | 0 | 0 |
| PAC1 | 2606217419 | LLI | n.d. | >99 | 1 | 1 | 1 | 1 |
| AG-402-P18 | 2667527282 | LLI | 0 | 94.66 | 1 | 1 | 0 | 0 |
| AG-402-O21 | 2667527279 | LLI | 1 | 94.38 | 1 | 0 | 1 | 1 |
| AG-402-I21 | 2667527261 | LLI | 0 | 92.53 | 1 | 1 | 0 | 0 |
| AG-402-B05 | 2716884375 | LLI | 1 | 92.26 | 1 | 1 | 1 | 1 |
| AG-402-B03 | 2716884374 | LLI | 1 | 91.3 | 1 | 1 | 1 | 1 |
| AG-402-I20 | 2667527260 | LLI | 0 | 90.81 | 1 | 1 | 0 | 0 |
| AG-402-L20 | 2667527270 | LLI | 1 | 89.92 | 1 | 0 | 1 | 1 |
| AG-402-A21 | 2667527256 | LLI | 0 | 89.8 | 1 | 1 | 0 | 0 |
| AG-402-M18 | 2667527273 | LLI | 0 | 89.43 | 1 | 1 | 0 | 0 |
| AG-402-I05 | 2667527259 | LLI | 1 | 87.77 | 0 | 0 | 0 | 0 |
| AG-402-B19 | 2716884732 | LLI | 1 | 87.41 | 0 | 0 | 1 | 1 |
| AG-402-N09 | 2716884384 | LLI | 1 | 87.36 | 1 | 0 | 1 | 1 |
| AG-402-L09 | 2667527268 | LLI | 1 | 87.09 | 1 | 1 | 1 | 1 |
| AG-402-C09 | 2716884377 | LLI | 1 | 86.91 | 1 | 0 | 1 | 1 |
| AG-409-A10 | 2716884385 | LLI | n.d. | 86.68 | 1 | 1 | 1 | 1 |
| AG-402-G08 | 2716884381 | LLI | 1 | 85.76 | 1 | 0 | 1 | 1 |
| AG-402-G06 | 2716884265 | LLI | 1 | 84.69 | 1 | 0 | 1 | 1 |
| AG-402-K14 | 2667527342 | LLI | 0 | 84.51 | 1 | 1 | 0 | 0 |
| AG-402-A08 | 2716884373 | LLI | 1 | 84.44 | 1 | 1 | 1 | 1 |
| AG-402-M23 | 2667527274 | LLI | 1 | 84.38 | 1 | 0 | 1 | 1 |
| AG-402-P16 | 2667527281 | LLI | 0 | 78.67 | 1 | 1 | 0 | 0 |
| AG-402-I22 | 2667527262 | LLI | 0 | 77.24 | 1 | 0 | 0 | 0 |
| AG-459-P07 | 2716884799 | LLI | 0 | 74.16 | 1 | 1 | 0 | 0 |
| AG-402-K04 | 2716884382 | LLI | 1 | 73.97 | 0 | 0 | 1 | 1 |
| AG-363-A03 | 2667527354 | LLI | 0 | 71.65 | 1 | 1 | 0 | 0 |
| AG-402-E17 | 2716884379 | LLI | 1 | 70.56 | 1 | 0 | 1 | 1 |
| AG-363-B05 | 2667527361 | LLI | 0 | 66.18 | 0 | 0 | 0 | 0 |
| AG-402-G19 | 2716884734 | LLI | 1 | 65.62 | 1 | 1 | 1 | 1 |
| AG-402-J18 | 2667527264 | LLI | 0 | 61.78 | 0 | 0 | 0 | 0 |
| AG-402-B10 | 2716884376 | LLI | 1 | 55.66 | 0 | 0 | 0 | 0 |
| AG-363-G03 | 2667527367 | LLI | 0 | 52.3 | 0 | 0 | 0 | 0 |
| AG-363-A04 | 2667527355 | LLI | 0 | 47.45 | 0 | 0 | 0 | 0 |
| AG-459-C18 | 2716884791 | LLI | 0 | 47.33 | 1 | 0 | 0 | 0 |
| AG-311-K21 | 2716884633 | LLI | n.d. | 46.35 | 1 | 0 | 1 | 0 |
| AG-363-M20 | 2667527376 | LLI | 1 | 45.64 | 1 | 1 | 1 | 1 |

| | | | | | | | | |
|---|---|---|---|---|---|---|---|---|
| AG-402-L19 | 2667527269 | LLI | 1 | 45.52 | 1 | 1 | 0 | 0 |
| AG-363-B19 | 2667527364 | LLI | 0 | 41.58 | 0 | 0 | 0 | 0 |
| MIT0602 | 2606217317 | LLII-III | n.d. | >99 | 0 | 0 | 0 | 0 |
| MIT0603 | 2606217686 | LLII-III | n.d. | >99 | 0 | 0 | 0 | 0 |
| MIT0918 | 2681812902 | LLII-III | n.d. | >99 | 1 | 1 | 0 | 0 |
| MIT1304 | 2681813571 | LLII-III | n.d. | >99 | 1 | 1 | 0 | 0 |
| SS120 | 2623620733 | LLII-III | n.d. | >99 | 0 | 0 | 0 | 0 |
| AG-402-G10 | 2716884733 | LLII-III | 0 | 91.3 | 1 | 1 | 0 | 0 |
| AG-363-B11 | 2667527362 | LLII-III | 0 | 82.05 | 1 | 1 | 0 | 0 |
| AG-363-C02 | 2667527365 | LLII-III | 0 | 77.85 | 0 | 0 | 0 | 0 |
| AG-363-I21 | 2667527369 | LLII-III | 0 | 68.49 | 1 | 1 | 0 | 0 |
| AG-363-G23 | 2667527368 | LLII-III | 0 | 33.97 | 1 | 1 | 0 | 0 |
| MIT0601 | 2606217319 | LLII-III | n.d. | >99 | 0 | 0 | 0 | 0 |
| MIT0919 | 2681812903 | LLII-III | n.d. | >99 | 0 | 0 | 0 | 0 |
| MIT9211 | 2623620960 | LLII-III | n.d. | >99 | 0 | 0 | 0 | 0 |
| AG-402-G22 | 2667527257 | LLII-III | 0 | 87.6 | 0 | 0 | 0 | 0 |
| AG-363-N16 | 2667527310 | LLII-III | 0 | 85.29 | 0 | 0 | 0 | 0 |
| AG-363-M21 | 2667527327 | LLII-III | 0 | 64.9 | 1 | 1 | 0 | 0 |
| AG-363-N03 | 2667527328 | LLII-III | 0 | 51.49 | 0 | 0 | 0 | 0 |
| MIT0701 | 2606217684 | LLIV | n.d. | >99 | 1 | 1 | 0 | 0 |
| MIT0702 | 2606217681 | LLIV | n.d. | >99 | 1 | 1 | 0 | 0 |
| MIT0703 | 2606217682 | LLIV | n.d. | >99 | 1 | 1 | 0 | 0 |
| MIT1205 | 2681813566 | LLIV | n.d. | >99 | 1 | 1 | 0 | 0 |
| MIT1227 | 2681813569 | LLIV | n.d. | >99 | 1 | 1 | 0 | 0 |
| MIT1303 | 2681812923 | LLIV | n.d. | >99 | 1 | 1 | 0 | 0 |
| MIT1306 | 2681812924 | LLIV | n.d. | >99 | 1 | 1 | 0 | 0 |
| MIT1312 | 2681812925 | LLIV | n.d. | >99 | 1 | 1 | 0 | 0 |
| MIT1313 | 2681812926 | LLIV | n.d. | >99 | 1 | 1 | 0 | 0 |
| MIT1318 | 2681812927 | LLIV | n.d. | >99 | 1 | 1 | 0 | 0 |
| MIT1320 | 2681812928 | LLIV | n.d. | >99 | 1 | 1 | 0 | 0 |
| MIT1323 | 2681812948 | LLIV | n.d. | >99 | 1 | 1 | 0 | 0 |
| MIT1327 | 2681812949 | LLIV | n.d. | >99 | 1 | 1 | 0 | 0 |
| MIT1342 | 2681812950 | LLIV | n.d. | >99 | 1 | 1 | 0 | 0 |
| MIT1418 | 2681813575 | LLIV | n.d. | >99 | 1 | 1 | 0 | 0 |
| MIT9303 | 2623620962 | LLIV | n.d. | >99 | 1 | 1 | 0 | 0 |
| MIT9313 | 2606217562 | LLIV | n.d. | >99 | 1 | 1 | 0 | 0 |
